# Ribosomal protein RPL39L is an efficiency factor in the cotranslational folding of proteins with alpha helical domains

**DOI:** 10.1101/2023.04.03.535332

**Authors:** Arka Banerjee, Meric Ataman, Maciej Jerzy Smialek, Debdatto Mookherjee, Julius Rabl, Aleksei Mironov, Lea Mues, Ludovic Enkler, Mairene Coto-Llerena, Alexander Schmidt, Daniel Boehringer, Salvatore Piscuoglio, Anne Spang, Nitish Mittal, Mihaela Zavolan

**Author notes:** These authors contributed equally (Meric Ataman, Maciej Jerzy Smialek).

## Abstract

Increasingly many studies reveal how ribosome composition can be tuned to optimally translate the transcriptome of individual cell types. In this study, we investigated the expression pattern, structure within the ribosome and effect on protein synthesis of the ribosomal protein paralog 39L (RPL39L). With a novel mass spectrometric approach we have quantified the expression of RPL39L in human pluripotent cells, cancer cell lines and tissue samples, and in mouse germ cells. We generated *RPL39L* knock-out mouse embryonic stem cell (mESC) lines and demonstrated that RPL39L impacts the dynamics of translation, to support the pluripotency and differentiation, spontaneous and along the germ cell lineage. Most differences in protein abundance between WT and RPL39L KO lines were explained by widespread proteasomal activity. By CryoEM analysis of purified RPL39 and RPL39L-containing ribosomes we found that, unlike RPL39, RPL39L has two distinct conformations in the exposed segment of the nascent peptide exit tunnel, creating a distinct hydrophobic patch that has been predicted to support the efficient co-translational folding of alpha helices. Our study shows that ribosomal protein paralogs provide switchable modular components that can tune translation to the protein production needs of individual cell types.

## Introduction

Protein synthesis is carried out by the ribosome, a highly conserved molecular machine with the same basic architecture in all free living organisms. In mammals, the small, 40S ribosomal subunit contains the 18S ribosomal RNA (rRNA) and 33 ribosomal proteins (RPs), while the large 60S subunit contains 46 RPs along with the 5S, 5.8S and 28S rRNAs. Gene duplications gave rise to RP paralogs ^1^, some with evolutionarily-conserved tissue-specific patterns of expression ^2^. Among these, autosomal-chromosome-encoded paralogs of X chromosome-linked RPs have strong expression bias for the male germ cell lineage, where they have been linked to highly specialized functions: *RPL10L* takes over the function of *RPL10* upon X chromosome inactivation in the meiotic phase of spermatogenesis ^3^, and RPL39L was implicated in the translation of long-lived, sperm cell-specific proteins ^4^. However, the *RPL39L* mRNA was also observed in other cell types, including ovarian ^5^ and breast cancers ^2^, as well as in lung cancer ^6^ and neuroblastoma ^4^ cell lines. Gene amplifications ^7^ and the hypomethylation of a specific CpG island ^6^ were proposed as mechanisms underlying RPL39L expression in cancers. These observations suggest that RPL39L has a general function, extending beyond the translation of long-lived sperm cell proteins. Unraveling this function has been challenging. RPL39L differs from RPL39 by only 4 or 3 amino acids in human and mouse, respectively, explaining why antibodies that can distinguish RPL39L from RPL39 are still lacking. In addition, the high arginine/lysine content of RPL39L leads to the almost complete digestion of the protein during standard sample preparation for mass spectrometry, probably restricting its detection to cell types with very high expression ^4^. As pure populations of RPL39L-containing ribosomes have not been obtained so far ^4^, how RPL39/RPL39L ribosomes differ has also remained unclear.

Our study aimed to determine the role of RPL39/RPL39L ribosome heterogeneity across mammalian cell types. We took advantage of mouse embryonic stem cells (mESC), a cell type with native expression of RPL39L, to generate RPL39L-deficient mESC lines by CRISPR/Cas9 genome editing. We then characterized their gene expression and capacity to differentiate both spontaneously and towards the sperm cell lineage. Ribosome footprinting along with mass spectrometric analysis in the presence and absence of protein degradation inhibitors revealed that RPL39L supports the translation of a heterogeneous collection of proteins. Many of these are involved in cell motility and polarization, thus explaining the critical role of RPL39L in spermatogenesis. As obtaining pure populations of RPL39L ribosomes from mice remains challenging, to analyze the impact of RPL39/RPL39L on ribosome structure we turned to the yeast, which has only the *RPL39* gene. We complemented *RPL39*-deficient yeast cells with either mouse *RPL39* or *RPL39L,* and analyzed purified RPL39 and RPL39L ribosomes. While mouse and yeast RPL39 occupy virtually identical positions in the ribosome exit tunnel, RPL39L was found in two distinct conformations, one very similar and the other distinct from RPL39. The alternative conformation creates a hydrophobic patch in the vestibular region of the nascent peptide exit tunnel (NPET), which was previously postulated to be necessary for the folding of amphipathic ɑ-helices ^8^. Our results provide an example of paralogous RPs supporting the generation of ribosomes with distinct biophysical properties. In particular, the flexibility conferred to the peptide exit tunnel of the ribosome by RPL39L relative to RPL39 appears to be important for the folding and stability of proteins with long helical domains, many of which are abundant in the male germ cells, but can be more generally described as relevant for cell motility and polarization.

## Results

### The RP paralog RPL39L is expressed in a variety of normal and malignant cells

Aiming to determine the breadth of *RPL39L* expression across normal human cells, we examined the single-cell sequencing (scRNA-seq) data from The Human Protein Atlas (HPA ^9^, https://www.proteinatlas.org/download/rna_single_cell_type_tissue.tsv.zip). We found that, while most abundant in the cells of the male germ cell lineage, the *RPL39L* mRNA is also present in other cell types, such as the extravillous trophoblast, where *RPL39L* level is ∼12-fold lower compared to spermatocytes (Fig. 1A, top). In contrast, another germ cell-specific RP, *RPL10L*, is virtually absent outside of the male germ line (Fig. 1A, top). For comparison, we also investigated the variation of mRNAs encoding core RPs (identified as described in Methods, Fig. S1) across cell types, finding it to be much smaller relative to the RP paralogs (Fig. 1A, top). The scRNA-seq data also gave us the opportunity to determine whether *RPL39L* replaces *RPL39* in specific cell types or rather the two genes are co-expressed. In the HPA scRNA-seq read count data (https://www.proteinatlas.org/download/rna_single_cell_read_count.zip) all cells that contained *RPL39L*-derived reads also had RPL39-derived reads, the sole exception being male germ cells, some of contained exclusively RPL39L reads. This suggests that some cell types have heterogeneous RPL39/RPL39L populations of ribosomes (Fig. 1A, bottom panel). The relatively high expression level of *RPL39L* in trophoblast cells (Fig. 1A. top) prompted us to further examine the dynamics of *RPL39L/RPL39* expression in early human development relative to adult human tissues. As the overall abundance of ribosomes differs across cell types ^2^, we always examined the variation in *RPL39L/RPL39* levels relative to the core RPs. Reanalyzing scRNA-seq data sets from pre-implantation embryos and embryonic stem cells (ESCs) ^10^ along with normal tissue samples in The Cancer Genome Atlas (https://www.cancer.gov/tcga/) we found that, relative to core RPs, *RPL39* maintained the same level in embryonic and differentiated cells (Fig. 1B), while the level of *RPL39L* was significantly higher in embryonic cells than in adult normal cells (Fig. 1B). The *RPL39L*-to-core RP ratio also varied more than 30-fold across cancer types (Fig. 1B), in many cancers reaching the values observed in embryonic cells. The ratio was highest in lung (LUSC) and cervix (CESC) tumors. In contrast, the *RPL39*-to-core RP ratio fluctuated much less across tumors (Ansari-Bradley test, p-value<10^-4^) and relative to normal cells.

**Figure 1.**
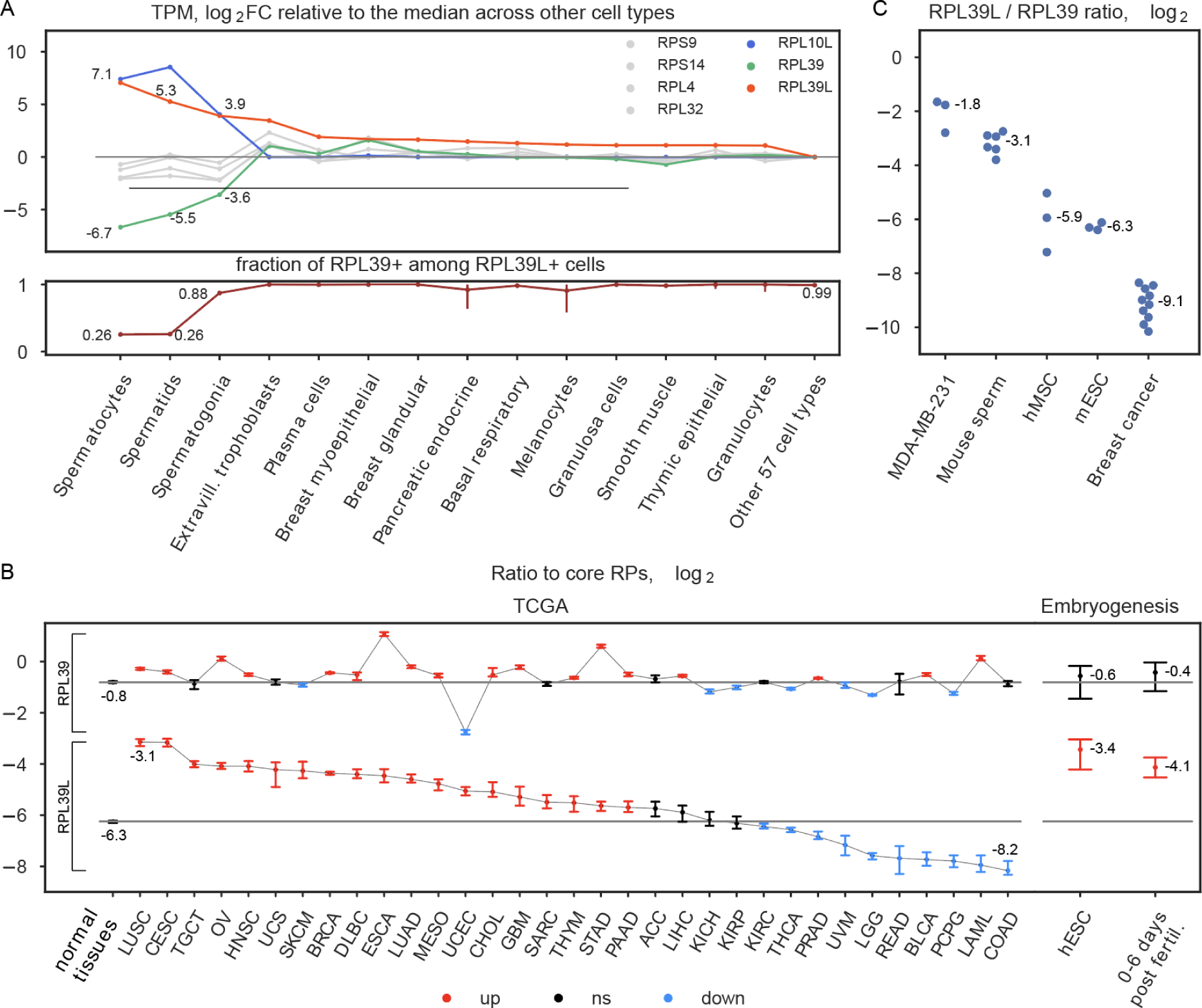
RPL39L expression across cell types. **A.** Top: HPA-provided normalized gene expression values (transcript-per-million, TPM) were used to identify the 14 cell types with highest *RPL39L* expression. The log_2_ fold change in each of these cell types relative to the median across all other 57 cell types in HPA is shown for *RPL39L* (orange), *RPL39* (green), *RPL10L* (blue) genes, and 4 core RPs (gray). Bottom: the proportion of *RPL39L*+ cells that also contained *RPL39*-derived reads in single cells of the types shown in the top panel. **B.** Ratio of *RPL39/RPL39L*-to-core RP expression (log_2_) per cell in human pre-implantation embryos and cultured embryonic stem cells ^10^ (left panel), as well as in bulk RNA-seq samples of primary tumors from TCGA (https://www.cancer.gov/tcga/) (right panel). See https://gdc.cancer.gov/resources-tcga-users/tcga-code-tables/tcga-study-abbreviations for TCGA cancer-type abbreviations. The median values of these ratios in normal samples are indicated by the black horizontal lines. Statistically significant (two-sided Wilcoxon test, Benjamini-Hochberg FDR<0.05) positive and negative deviations from the medians are shown in orange and blue, respectively. For each sample type, the 95% confidence interval over all samples is shown. Sample types for which the ratios were not significantly different from the median of the normal samples are shown in black. **C.** Quantification of RPL39L/RPL39 protein ratio in various cellular systems (mouse sperm cells - 6 samples, breast cancer cell line MDA-MB-231, bone marrow-derived mesenchymal stem cells (hMSC) and mouse embryonic stem cells (mESC) - 3 independent samples, breast cancer tissues - 10 samples) using reference peptides.

To validate the mRNA level data, we sought to measure the RPL39L protein in a few cellular systems. As RPL39L-specific antibodies are not available, we turned to mass spectrometry. Due to its high lysine/arginine content, the RPL39L protein is not reliably captured in standard proteomics analyses. Thus, we modified the sample preparation, acetylating the lysines in the proteome to direct the cleavage by trypsin to arginines only. We used heavy-labeled reference peptides to measure the RPL39L-to-RPL39 protein ratio in mature mouse sperm cells as positive control, and then in mouse embryonic stem cells, the human MDA-MB-231 breast cancer cell line, as well as human tissue samples, namely bone marrow-derived mesenchymal stem cells and breast cancer tissues (Fig. 1C). The values indicate that, if the measured proteins come primarily from ribosomes, ∼30% of ribosomes in MDA-MB-231 cell line, ∼10% of ribosomes in mature mouse sperm cells, and ∼1% of ribosomes in bone marrow stromal cells and mouse embryonic stem cells contain RPL39L.

In human breast cancer tissue samples with heterogeneous cell type composition, RPL39L is still detectable, though at a much lower abundance (Fig. 1C). Our data thus provide conclusive mass spectrometric evidence of RPL39L protein expression not only in male germ cells, but also in pluripotent cells and cancer cell lines. The broader expression pattern of RPL39L compared to other germ cell-biased RP paralogs points to the relevance of RPL39/RPL39L ribosome heterogeneity beyond spermatogenesis.

### KO of *RPL39L* impairs the pluripotency of E14 mESCs

To determine the role of RPL39L in pluripotent cells we generated *RPL39L* knockout (KO) mESCs by CRISPR-mediated deletion of the coding region of this gene in the E14 mouse stem cell line. To minimize the chance of interfering with *RPL39* we designed the sgRNAs to target non-coding regions of *RPL39L*, which are not shared with *RPL39*. To further exclude sgRNA-specific off-target effects, we designed two sets of guide RNAs that matched non-coding regions around the *RPL39L* CDS (Fig. 2A), and for each sgRNA set we selected clones that originated from independent editing events (Fig. S2). We comprehensively analyzed two distinct clones for each sgRNA pair, in which the editing of the *RPL39L* locus was validated both by qPCR amplification with probes that flanked the edited region, and by measuring the protein expression with targeted proteomics (Fig. 2B-C). All four clones (labeled as 1.17, 1.20, 2.9 and 2.11) exhibited homozygous deletions of the *RPL39L* locus, resulting from a spectrum of editing events including seemingly distinct deletions on the two chromosomes in clone 2.11 (Fig. S2). The RPL39L protein was undetectable in all of the edited clones except for 1.17, where some residual protein expression was detected.

**Figure 2.**
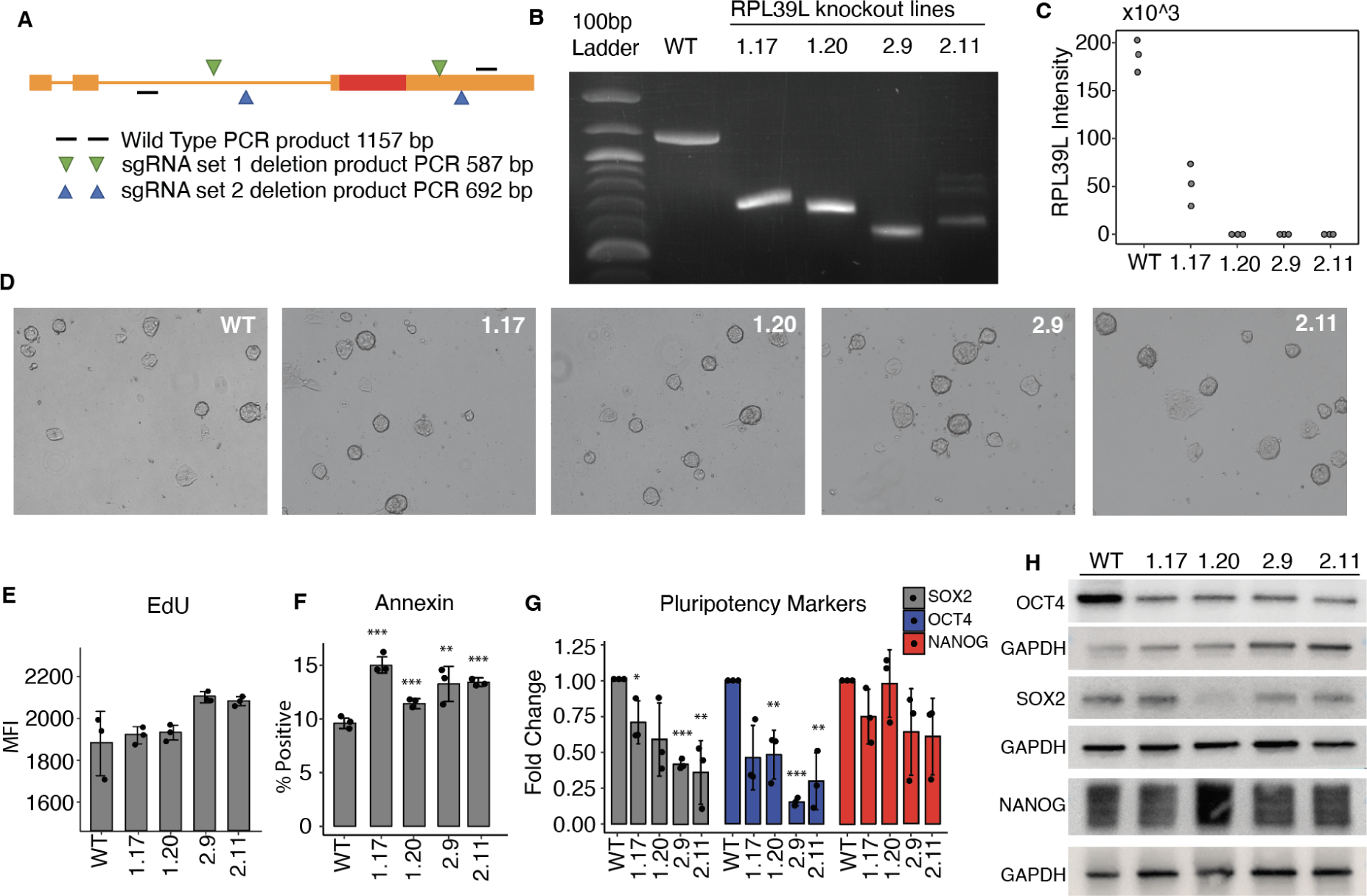
Characterization of *RPL39L* KO mESCs. **A.** Schema of sgRNA design. Two distinct pairs of sgRNAs (in green and blue) were designed to target flanking regions of the *RPL39L* coding sequence (CDS, shown as a red box). Primers (black lines) from further upstream and downstream in the *RPL39L* locus were used for amplification, and the expected sizes of the PCR products are indicated for both the WT locus (1157 nts) and the edited loci (587 and 692 nts, respectively, for the sgRNA sets 1 and 2). **B.** PCR products from the WT E14 cells and the 4 independent clones, 1.17 and 1.20 generated with sgRNA set #1 and2.9 and 2.11 generated with sgRNA set #2. **C.** Expression of RPL39L in individual clones determined with targeted proteomics (n=3 for each clone). **D.** Representative bright field images of colonies from all analyzed clones. **E.** Results of the ethynyldeoxyuridine (EdU, thymidine analog) incorporation assay. EdU-treated cells were fixed, permeabilized and the AF488 fluorophore was linked to the EdU in the replicated DNA by click-chemistry. The label intensity was measured by FACS (Mean fluorescence intensity, MFI). **F.** Results of the annexin binding assay. Cells were incubated with AF488-conjugated Annexin V and counterstained with phycoerythrin (PE). The proportion of AF488^+^PE^-^ (apoptotic cells) relative to the parent cell population was determined by FACS. **G.** RT-qPCR of the pluripotency factors *SOX2*, *OCT4* and *NANOG*. Values are 2^-ΔΔCt^, relative to *RRM2* (internal reference) and to WT. **H.** Representative Western blots of pluripotency markers SOX2, OCT4 and NANOG in the RPL39L KO lines and WT relative to GAPDH. In panels F and G *, ** and *** correspond to p-values <0.05, <0.01 and <0.001, respectively in the two tailed t-test comparing KO lines with WT.

The *RPL39L* KO cells were viable in culture, forming colonies that were similar in morphology to those formed by the WT mESCs (Fig. 2D). Analysis of 5-ethynyl-2-deoxiuridine (EdU) incorporation during 2 hours of treatment did not reveal significant differences between KO and WT cells, indicating that the *RPL39L* KO does not significantly impact proliferation in mESCs (Fig. 2E). Annexin V staining showed only small, though statistically significant increase in the proportion of apoptotic cells relative to WT (Fig. 2F). Despite no obvious changes in the colony morphology, the pluripotency markers exhibited significant variation; the Sox2 and Oct4 levels were reduced in KO compared to WT E14 lines, while Nanog showed less consistent reduction (Fig. 2G). These differences were also apparent at the protein level (Fig. 2H). Thus, *RPL39L* KO lines are viable, but have reduced expression of pluripotency markers relative to WT mESCs.

### *RPL39L* KO mESCs exhibit differentiation defects

The perturbed expression of pluripotency markers in *RPL39L* KO cells prompted us to further investigate their ability to differentiate. Given the relevance of RPL39L for the spermatogenic lineage ^4, 11^, we first subjected WT and *RPL39L* KO E14 lines to *in vitro* differentiation along this lineage using a previously described protocol ^12^. The emergence of spermatocyte-like cells in the WT E14 mESC culture demonstrated that the protocol works as expected (Fig. 3A). In contrast, we did not identify any spermatocyte-like cells in the cultures of *RPL39L* KO lines, and the expression of germ-cell lineage markers Stella and Dazl were significantly reduced relative to differentiating WT cells (Fig. 3B). These results show that RPL39L is necessary for spermatogenesis *in vitro*, consistent with previous observations in mice ^4, 11^. To determine whether the KO of *RPL39L* impacts other developmental lineages as well, we carried out spontaneous differentiation of all our mESC lines *in vitro*, by culturing the cells in leukemia inhibiting factor (LIF)-free medium under non-adherent conditions ^13^. We then analyzed the expression of lineage markers both by qRT-PCR and by immunofluorescence-based quantification of protein levels in embryoid body sections (Fig. 3C-F). The qRT-PCR revealed decreased expression of extraembryonic endoderm markers *GATA6, GATA4 and DAB2* in the organoids generated from the KO cells relative to those generated from WT cells (Fig. 3C,E) and increased expression of ectodermal lineage markers *NESTIN, FGF5 and PAX6* (Fig. 3D,E). The protein level quantification from confocal images of NESTIN (ectoderm) and GATA4 (endoderm) validated the mRNA-level results (Fig. 3F). Thus, the KO of *RPL39L* impacts spontaneous differentiation of mESCs, primarily towards the ectoderm and endoderm.

**Figure 3.**
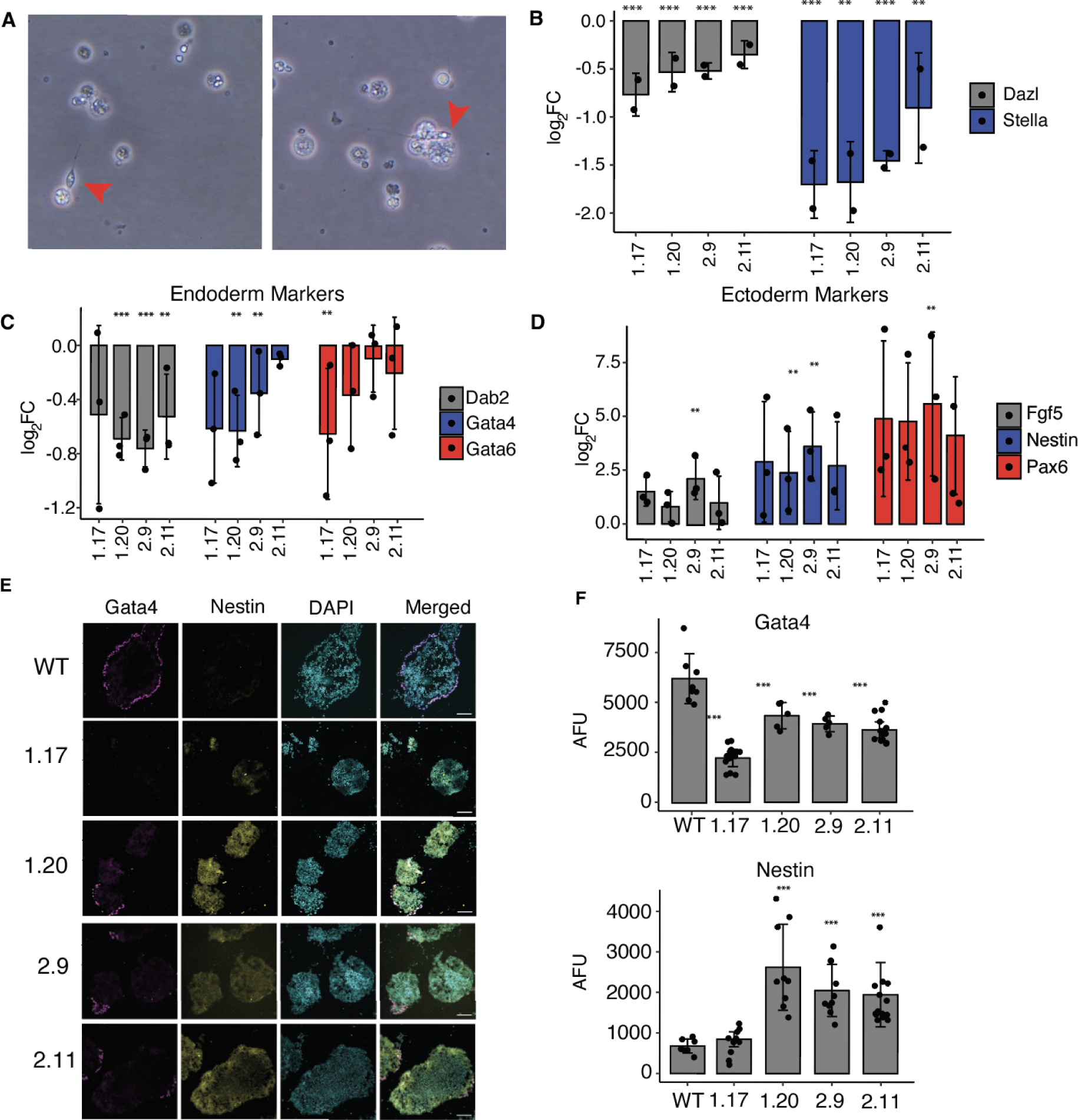
*RPL39L* KO leads to differentiation defects in E14 mESCs. **A.** Representative bright field images showing the spermatogenic differentiation of WT E14 cells. Red arrows indicate spermatocyte-like cells. **B.** qRT-PCR assays of *DAZL* (late) and *STELLA* (early) sperm cell markers ^12^ (y-axis, log_2_ fold-change) in differentiating (4 days in RA-containing medium) populations of KO clones (x-axis) relative to WT. **C.** qRT-PCR of extraembryonic endoderm markers (*DAB2, GATA4, GATA6*) ^13^ in *RPL39L* KO lines relative to WT. **D.** Similar for *FGF5, NESTIN*, and *PAX6* ectoderm markers ^13^. **E**. Immunofluorescence staining of embryoid bodies subjected to spontaneous differentiation: GATA4 was used as endoderm marker, NESTIN as ectoderm marker and DAPI to delineate the nucleus. **F.** Quantification of GATA4 and NESTIN expression in immunofluorescence images. AFU - arbitrary fluorescence units normalized to DAPI. In all panels, *, ** and *** correspond to p-values <0.05, <0.01 and <0.001, respectively in the two tailed t-test comparing KO lines with WT.

### *RPL39L* KO lines exhibit perturbed protein synthesis and ER stress

RPL39L being a ribosome component, to unravel the mechanisms underlying the observed functional defects, we evaluated the effect of *RPL39L* KO on global translation. We generated polysome profiles from all clones, and found that the KO of *RPL39L* leads to a small, but statistically significant increase in polysome-to-monosome ratio (Fig. 4A-B). This indicates that *RPL39L* KO induces either a small increase in the rate of translation ^14^ or to an elongation defect ^15^ in mESCs.

**Figure 4.**
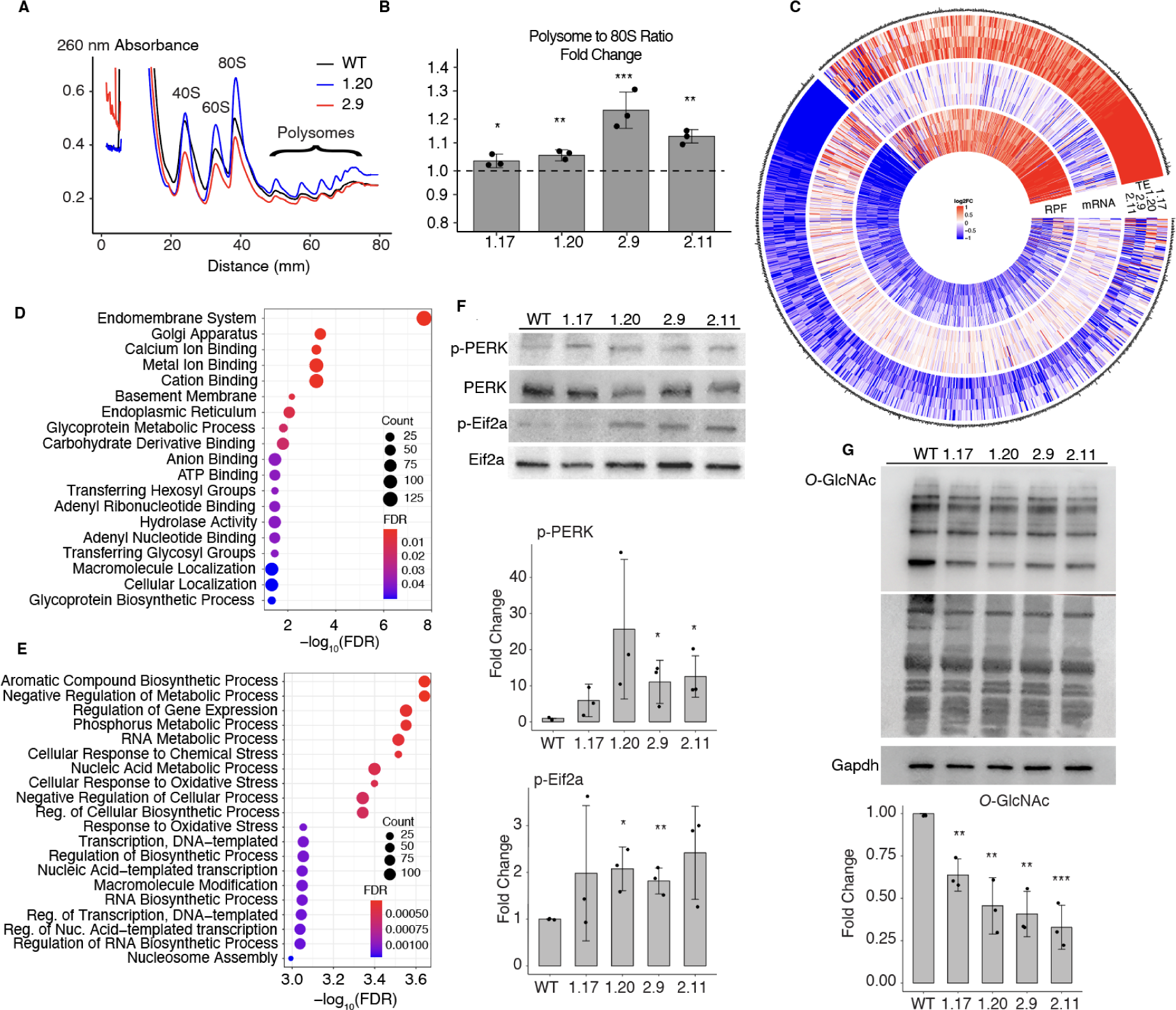
Impact of *RPL39L* KO on mRNA translation. **A.** Example polysome profiles from the WT, 1.20 and 2.9 RPL39L KO E14 cell lines. **B.** Ratio of the area under the profile corresponding to polysomes vs. monosomes (80S), in polysome profiles obtained from the KO clones. Fold-changes were calculated relative to the median ratio in the corresponding WT (dashed line at 1, n=3 for all cell lines). **C.** Log_2_ fold-changes in the translation efficiency (TE), mRNA level and the number of ribosome protected fragments (RPF) for specific genes, in mutant clones relative to WT E14 cells (n=3 for each clone). Shown are all genes with a significant change in TE in at least one of the *RPL39L* KO clones. Values are capped at −1 and 1. **D-E.** Gene Ontology analysis of mRNAs with reduced (D) and increased (E) TE in *RPL39L* KO clones. **F.** Representative western blots and corresponding quantification (from n=3 for each clone) of UPR markers PERK (phospho-Thr980) and EIF2A (phospho-Ser51). Intensities of phosphorylated proteins were normalized by the respective unphosphorylated forms and are relative to WT, for which the relative phosphorylation level was set to 1. **G.** Representative western blot and corresponding quantification (from n=3 for each clone) showing lower global O-GlcNAc modification of proteins in *RPL39L* KO lines when compared to the WT. Values are relative to GAPDH (loading control) and WT (level set to 1). In all panels, *, ** and *** correspond to p-values <0.05, <0.01 and <0.001, respectively in the two tailed t-test comparing KO lines with WT.

To determine whether some transcripts are specifically impacted in translation by the *RPL39L* KO, we carried out ribosome footprinting ^16^, sequencing ribosome-protected mRNA fragments (RPFs) from both WT and *RPL39L* KO mESCs. The RPF data fulfilled expected quality criteria such as the vast majority of reads mapping to the coding regions of mRNAs and the 3 nucleotide periodicity of inferred P site locations (Fig. S3). We further sequenced the mRNAs from these cells and calculated the translation efficiency (TE) per mRNA as the ratio of the RPF and mRNA read density along the CDS (see Methods). Focusing on mRNAs whose TE was significantly altered (p-value < 0.01 in ΔTE test, see Methods) in at least one of the KO lines relative to the WT, we found that while the differences in RPFs and TE in KO clones relative to WT were generally small, the direction of change was highly consistent among specific classes of mRNAs (Fig. 4C). Gene Ontology analysis showed that transcripts encoding components of the endomembrane system, including the Golgi apparatus and the endoplasmic reticulum (ER) experienced the strongest reduction in TE across the KO lines (Fig. 4D). In contrast, we found a significantly increased TE for transcripts associated with the cellular response to chemical and oxidative stress (Fig. 4E). Examples of increased and decreased RPF coverage of specific genes are shown in Fig. S3. These results suggest that the production of specific classes of proteins, associated with subcellular compartments such as the ER and Golgi apparatus, is impaired in *RPL39L* KO cells.

To further elucidate the relationship between the *RPL39L* KO and cellular stress, we measured the levels of both phosphorylated PKR-like ER kinase (PERK), a sensor of ER stress ^17^ and of its phosphorylation target, the eukaryotic initiation factor 2a (EIF2A) ^18^, which is responsible for regulation of many unfolded protein response (UPR)-associated stress-response genes. Western blotting showed that both of these markers were elevated in *RPL39L* KO compared to WT E14 cells (Fig. 4F). We further used an O-GlcNAc antibody to label the glycosylated proteins from all our cell lines on a western blot. Integrating the chemiluminescence signal over the entire lanes corresponding to specific cell lines, we found reduced global signals in the samples from *RPL39L* KO cell lines compared to the WT E14 line (Fig. 4G). Thus, ER stress markers are upregulated and protein glycosylation is impaired in *RPL39L* KO mESC lines relative to WT.

### Increased degradation underlies the perturbed protein levels in *RPL39L* KO cell lines

To learn more about the protein synthesis in *RPL39L* KO and WT cells, we measured the protein levels, both in steady-state and upon inhibition of proteasome and autophagy-dependent protein degradation (by treatment with MG132 and Bafilomycin A1, respectively), by shot-gun mass spectrometry. Overall, we identified 5’026 proteins, 2’992 of which in both treated and untreated conditions (Fig. S4). Strikingly, many proteins whose expression was perturbed by the *RPL39L* KO were restored to levels similar to those in WT by the inhibition of protein degradation, as indicated by the narrower distribution of log_2_ fold-changes of KO relative to WT cells in protease inhibitor-treated compared to untreated cells (Fig. 5A). Along with the observed ER stress, this points to a reduced stability of proteins in *RPL39L KO* lines, which in turn suggests an increased production of defective proteins. This could be due to translation errors, e.g. amino acid misincorporation, or to defects in co-translational protein folding. The proteins whose reduced level in a KO clone relative to WT was restored by the treatment with protease inhibitors are shown in Fig. 5B, which demonstrates their consistent behavior across KO clones. Western blotting further confirmed these results (Fig. 5C-D). Gene Ontology analysis revealed enrichments in categories associated with the cytoskeleton and microtubules, indicating that the proteins stabilized by the protease inhibitor treatment are components of cellular membranes, contributing to cytoskeletal organization (Fig. S4). With a previously developed tool that specifically searches for amino acid substitutions in the measured peptides ^19^ we were unable to detect an enrichment of misincorporation in these proteins (Fig. S4). Thus, the most plausible explanation for the observed proteome changes is that RPL39L ribosomes facilitate the co-translational folding of specific proteins, leading to the production of defective proteins with compromised stability in the *RPL39L* KO clones.

**Figure 5.**
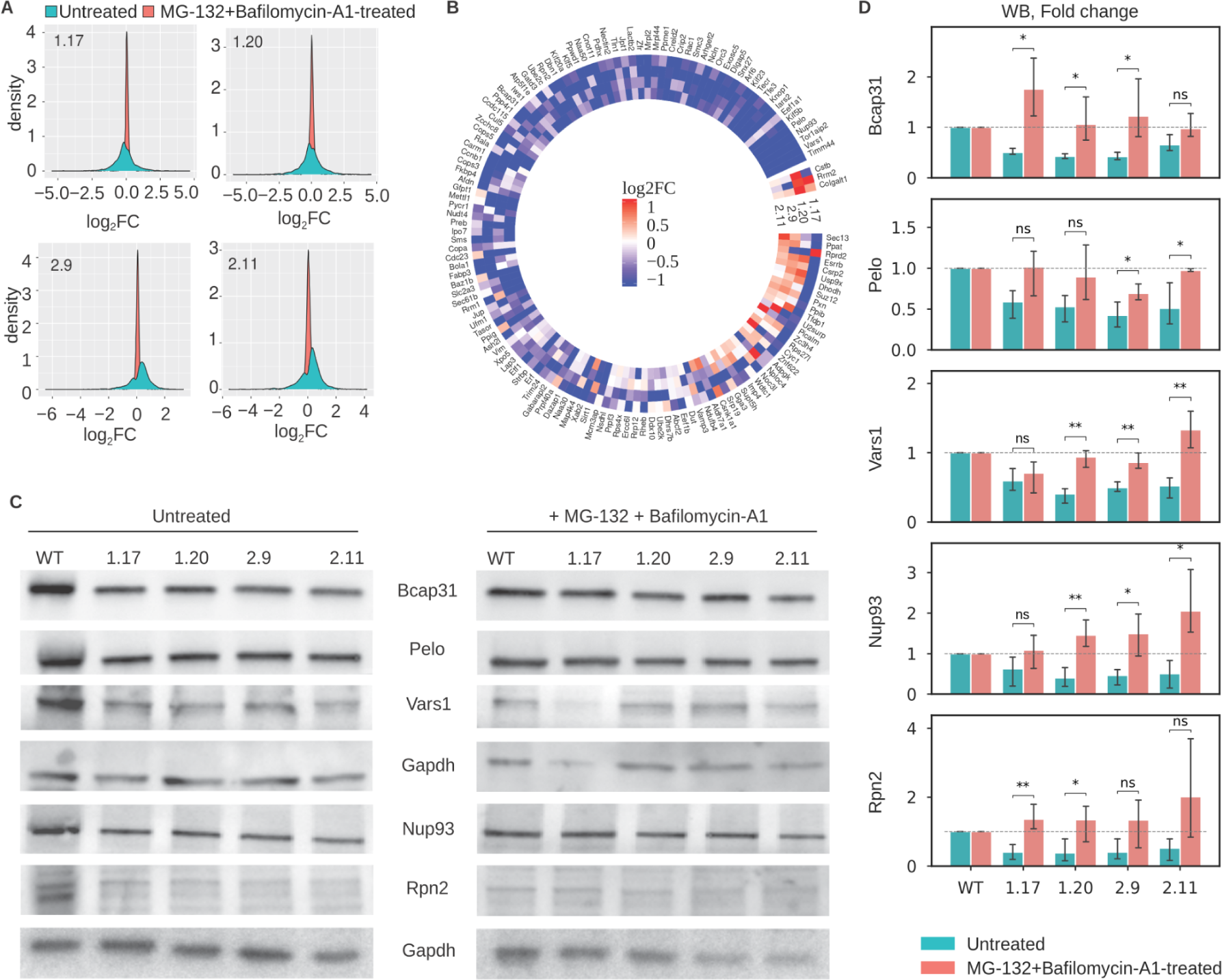
*RPL39L* KO clones exhibit enhanced degradation of specific classes of proteins. **A.** Distribution of log_2_ fold changes in protein levels between untreated (blue) or MG-132+Bafilomycin-A1-treated *RPL39L* KO and WT cells. Each panel corresponds to one KO clone. 3 biological replicates for each condition were used to calculate average protein abundance levels and respective fold-changes relative to WT. **B.** Heatmap of protein-level log_2_ fold changes in KO cells relative to WT. Included are all proteins with a significant downregulation restored by the protease inhibitor treatment in at least one of the KO clones. **C.** Representative western blot results showing the expression of a subset of proteins from (B) in untreated (left) and MG-132+Bafilomycin-A1-treated cells (right). **D.** Quantification of western blots as shown in (B), from three replicates for each protein and each condition.

### Conformational differences between RPL39/RPL39L ribosome peptide exit tunnels

To understand how RPL39L could influence the dynamics of translation and the co-translational folding of proteins we sought to analyze RPL39L and RPL39-containing ribosomes by cryo-electron microscopy (cryo-EM). Mouse RPL39L has only 3 substitutions relative to RPL39: S2A, R28Q and R36M. Since these differences are subtle, structural investigations of RPL39/RPL39L-ribosomes necessitate high-resolution structures, which can only be obtained from homogeneous (RPL39 or RPL39L only) samples. Obtaining pure populations of RPL39L-containing ribosomes has been an unsolved challenge ^4^, which we have decided to overcome by expressing the mouse RPL39 and RPL39L in yeast *RPL39* KO cell lines (Fig. 6A). In yeast, just like in mammals, residues R28 and R36 of RPL39 face the lumen of the NPET and are in direct proximity to the nascent polypeptide chain.

**Figure 6.**
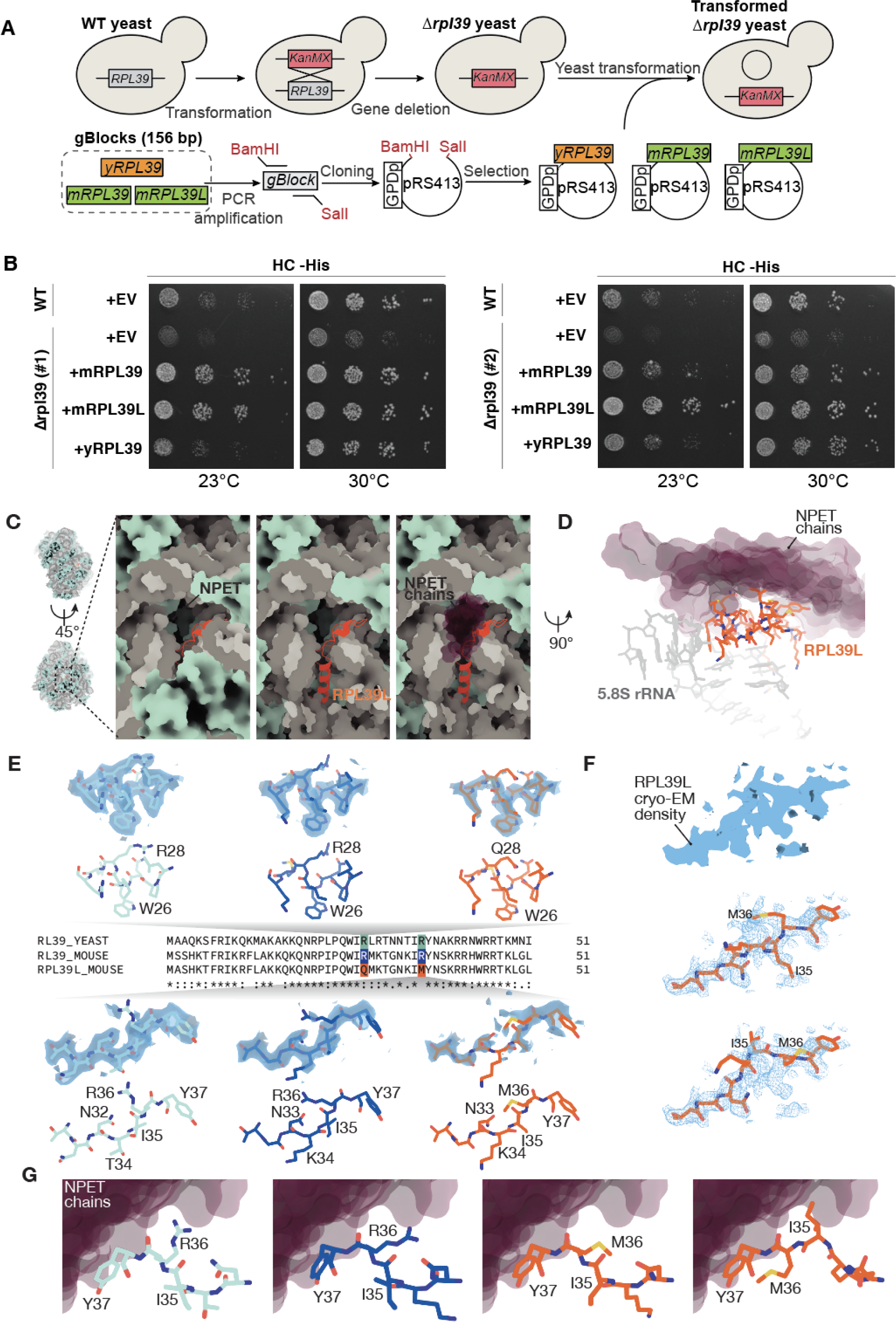
RPL39L introduces a hydrophobic patch in the NPET. **A.** Scheme of yeast RPL39 KO and insertion of mouse RPL39 and mouse RPL39L. **B.** The *RPL39* KO causes growth defects in yeast upon environmental challenges like growth at 23 degrees Celsius in HC-His media. Expression of mouse RPL39 or RPL39L rescues these phenotypes. +EV are cells transformed with an empty vector as control. The experiments were carried out in two independent RPL39 knockout clones (#1 and #2) **C.** RPL39L (cartoon representation, orange) shown in context of the large ribosomal subunit (refined atomic model of rpl39Δ-MmRpl39l; rRNAs are shown in gray, proteins are shown in pale blue surface representation). RPL39L is embedded inside the 60S subunit, adjacent to the exit of the NPET. Part of the 5.8S rRNA and ribosomal proteins have been removed for clarity on the center and right panel. Nascent protein chains and regulatory protein complexes pass the NPET in direct proximity to RPL39L as evident from aligned structures containing NPET-bound chains (nascent chains and regulatory proteins, PDB-IDs: 6M62, 7OBR, 7TM3, 7TUT, 7QWQ, 7QWS, shown as purple semi-transparent surfaces). **D.** Side view of RPL39L (orange) and the surrounding 5.8S rRNA (gray) in direct contact with the NPET-facing region of RPL39L. The region containing Q28 and M36 is located directly adjacent to protein chains localized in the lumen of the NPET in structures containing nascent chains or tunnel-bound regulatory complexes (protein chain models shown as purple semi-transparent surfaces). **E.** Comparison of the atomic model in the immediate surrounding of R/Q28 and R/M36 in WT yeast RPL39 (cyan), mouse RPL39 (dark blue), and mouse RPL39L (orange), shown in stick representation. The experimental cryo-EM density (semi-transparent surface, light blue) is shown superimposed on the refined atomic model. Maps around R28 are shown at a threshold of 6.5σ (yRPL39), 5.5σ (mRPL39), and 4.25σ (mRPL39L), while maps around R/M36 at a threshold of 5σ (yRPL39), 4σ (mRPL39), and 3.75σ (mRPL39L). Experimental cryo-EM density around M36 in mRPL39L is substantially weaker than the density observed either in yeast or mouse RPL39, due to increased conformational heterogeneity. **F.** At a lower threshold, an alternative conformation is apparent in the experimental cryo-EM density (blue surface) of the region around M36 in mRPL39L, as RPL39L adopts two alternative conformations that differ substantially relative to the protein backbone and side chains. **G.** In both WT yRPL39 (atomic model, stick representation, cyan) and mRPL39 (dark blue), the side chain of R36 faces the lumen of the NPET, potentially in direct contact with the protein chains inside the NPET (fitted chains of nascent protein and regulatory complexes, shown as purple semi-transparent surfaces). In mRPL39L, side chains of M36 and I35 hydrophobic residues are facing the NPET chains, forming a hydrophobic spot inside the NPET.

Thus, the replacement of R28 by Q28 and R36 by M36 in RPL39L, that is, of positively-charged residues by polar and hydrophobic residues, could perturb the co-translational folding or localization of proteins. The high degree of structural conservation of the 60S ribosomal subunit core in general, and of the region directly surrounding RPL39 in particular (mouse: RMSD=0.52 Å, human: RMSD=0.25 Å) further justifies the use of a heterologous system ^20^.

Increased rate of translation error reduces the growth of *RPL39* KO (*rpl39*Δ) yeast strains at cold temperatures and sensitizes the cells to translation-interfering drugs like paromomycin ^21^ and azetidine-2-carboxylic acid (AZT) ^22^. We replicated these phenotypes in two distinct yeast *RPL39* KO clones and further showed that they are rescued by the introduction of yRPL39 (rpl39Δ-ScRPL39 clones), mouse RPL39 (rpl39Δ-MmRPL39 clones) and mouse RPL39L (rpl39Δ-MmRPL39L clones) (Fig. 6B and S5). This indicates that both mRPL39 and mRPL39L occupy the place of yRPL39 in the NPET and have a similar capacity to ensure the translation accuracy ^21^ and promote co-translational folding ^22^ in yeast. The results also suggest RPL39L shares the ancestral function of RPL39, though it may have acquired additional functions following its emergence in mammalian species.

Ribosomal particles from the WT yeast strain Sc60S, the yeast *RPL39* KO strain rpl39Δ, yeast rescue strain rpl39Δ-ScRPL39 and the mouse RPL39/RPL39L-complemented *RPL39* KO strains rpl39Δ-MmRPL39 and rpl39Δ-MmRPL39L were studied by single-particle cryo-EM (Fig. S6, S7). The structure of the rpl39Δ ribosome demonstrated the absence of *RPL39* in the KO strain, with an otherwise structurally unaltered 60S subunit (overall RMSD = 0.308 Å), including the region surrounding *RPL*39 (RMSD=0.38 Å) (Fig. S7). Expression of yRPL39 in a RPL39Δ background resulted in a correctly assembled 60S subunit that did not differ significantly structurally from the wild type 60S (RMSD=0.192 Å), corroborating the feasibility of the complementation approach (Fig. S7). Mouse RPL39 and RPL39L both integrate into yeast 60S subunits when expressed in the rpl39Δ background (Fig. S7). Mouse RPL39, adopted a structure that was highly similar to that of RPL39 (RMSD=0.482 Å) in the WT yeast ribosomes (Fig. S7). The structure of RPL39 in mouse (PDB: 6SWA, RMSD=0.546 Å, ref. ^23^) and in human ribosomes (PDB: 7OW7 ^24^, RMSD=0.451 Å) is also in agreement with the structure of rpl39Δ-MmRPL39, suggesting that our heterologous system adequately reflects the structure and conformation of mammalian RPL39 (Figure S7). The comparison of rpl39Δ-MmRPL39 and rpl39Δ-MmRPL39L maps shows that the amino acids S2 in RPL39 and A2 in RPL39L are found at the same position, deeply embedded in rRNA 1490-1493, where they serve a structural role. The loop containing R28 and R36, exposed to the lumen of the NPET (Fig. 6C-E), exhibits weaker cryo-EM density than the remainder of the protein in the map of rpl39Δ-MmRPL39. The exchange of R28 and R36 with glutamine and methionine, respectively, in RPL39L further increases the flexibility (Fig. 6D-E) and enables the loop (residues 32-37) to adopt an alternative conformation where the Cɑ atom of I35 is displaced by 5.2 Å towards the exit of the NPET (Fig. 6E-F). The positions of the basic R28 side chain in RPL39 and polar Q28 side chain in RPL39L are nearly identical. While residues M29, K30, T31, and G32 adopt the same conformation in RPL39 and RPL39L, the side chain of N33 has a slight displacement of Cγ: 1.87 Å. The side chain of K34, which is located in direct proximity to the phosphate backbone of A351 in RPL39, protrudes into the lumen of the tunnel in RPL39L. However, a partial occlusion of the tunnel appears unlikely as the side-chain cryo-EM density for K34 is very weak. The position of I35, which is facing towards the wall of the tunnel (A351 and A42) in RPL39, is occupied by M36 in RPL39L, leaving I35 oriented towards the lumen of the tunnel in RPL39L in the alternative conformation (Fig. 6G). These observed structural changes do not substantially alter the overall electrostatic surface potential of the tunnel-exposed surface of RPL39/RPL39L, which is overwhelmingly positive in both cases. However, they do affect a narrow part of the ribosomal tunnel, traversed by the nascent chain (Fig 6D,E). Specifically, in RPL39L, Q28 and M36/I35 are found directly adjacent to each other at the location occupied by R28 and R36 in RPL39, introducing a clearly defined hydrophobic patch at the surface of the tunnel (Fig. 6G). This may influence the translation speed, efficiency or the co-translational folding, either via direct interaction with the nascent chain (Fig. 6G) or via interaction with translation regulation machinery such as the NAC complex, which was shown to bind to RPL39 ^25^.

### Altered translation dynamics of *RPL39L* KO-destabilized proteins

To determine how these conformational changes in the NPET may impact co-translational processes, we returned to the ribosome footprinting data, as such data were previously used to investigate multiple aspects of protein synthesis, including co-translational folding (reviewed in ^26^). We estimated the change in ribosome dwell time on individual codons between WT and *RPL39L* KO clones as described before ^27^ and then calculated the average changes over all codons encoding a particular amino acid. We found that histidine-encoding codons had consistently decreased ribosome dwell times in all of the KO lines relative to WT, while leucine-encoding codons had increased dwell time (Fig. 7A). For other amino acids the changes were smaller and less consistent between clones (Fig. 7A). Histidine and leucine are amino acids that are enriched in helix-turn-helix, zinc finger, leucine zipper coiled-coil motifs, which occur in proteins of diverse functions ^28–30^. Among the 39 proteins that were consistently destabilized in the KO lines, 7 contained coiled-coils. Interestingly, 6 of these (SMC3, KIF23, KIF5B, KIF20A, DLGAP5, BCAP31) have spermatocytes and/or trophoblast as the tissues of highest expression in the HPA (Fig. 7B). Along with the ribosome profiling and protein stability data, these data indicate that RPL39L ribosomes support the co-translational folding and stability of proteins with His/Leu-containing domains.

**Figure 7.**
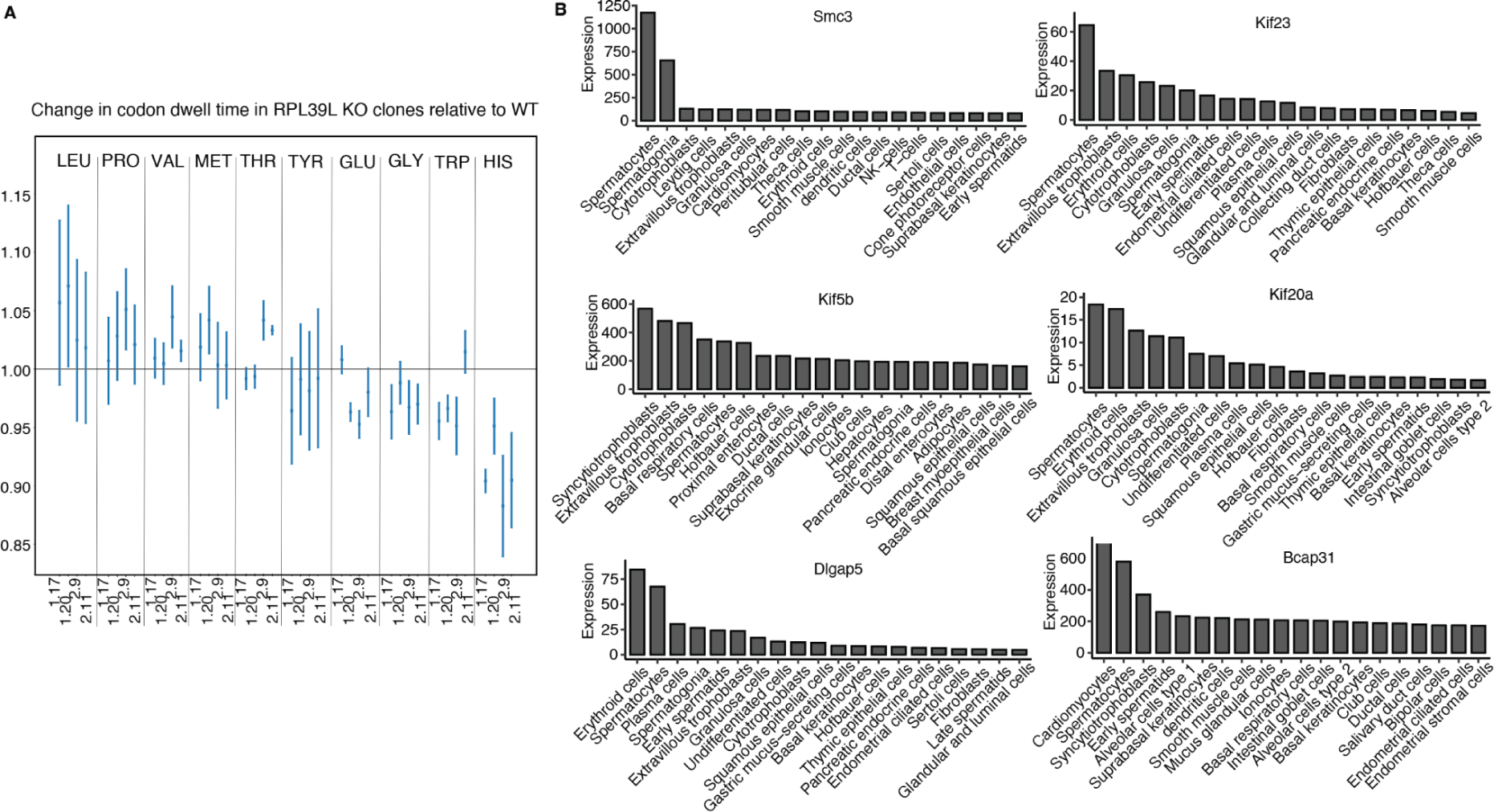
Altered codon dwell times in *RPL39l* KO relative to WT clones. **A.** Relative change in codon dwell time for codons encoding individual amino acids in the *RPL39L* KO clones relative to WT. The top five amino acids with the most increased and decreased average dwell time per codon are shown, and labeled at the top of the panel. Error bars over 3 replicates of each clone (indicated by x-axis tick labels) are shown. The black line drawn at 1 indicates no change in dwell time relative to the WT clone. **B.** Expression level (TPM) of mRNAs encoding proteins with coiled-coil domains that are destabilized in the *RPL39L* KO clones. Shown are values obtained by scRNA-seq of various cell types in the Human Protein Atlas. The gene names are indicated at the top of each panel and the cell types are labeled on the x-axis.

## Discussion

mRNA translation is carried out by the ribosome, a highly conserved molecular machine dating back to the last universal common ancestor of cellular life ^31^. Although a ribosome can, in principle, translate any of the mRNAs expressed in a cell, the protein output per mRNA, dependent on the efficiency of translation initiation, the speed of peptide chain elongation ^32, 33^, and factors such as the total number of ribosomes in the cell ^34–36^. In recent years, cell type-specific translational programs controlled by cell type-specific ribosomal protein paralogs ^37^ or by post-translational modifications of ribosome-associated proteins ^38^ have been described. The human and mouse genomes encode around 20 RP paralogs, which have a more restricted distribution across tissues than canonical RPs ^2^. RPL39L is a recently evolved RP paralog ^1^ recently found to be critical to male fertility ^4, 11^. The *RPL39L* mRNA is also expressed outside of the male germ line, in normal and cancer cells ^2, 5, 6^, where its role is still unknown. Furthermore, while RPL39L has been implicated in the folding of long-lived sperm proteins ^4, 11^, how this is achieved is unclear.

To answer these questions, we generated *RPL39L* mESC KO lines by CRISPR-mediated genome editing. ESCs naturally express *RPL39L*, as we have found in analyses of bulk and scRNA-seq datasets. To confirm *RPL39L* expression at the protein level, we developed a mass spectrometric-based approach wherein lysine residues are acetylated proteome-wide to direct the tryptic cleavage to arginine residues. With its high content of lysines and arginines (31% of 51 amino acids), RPL39L is almost completely digested by trypsin during standard sample preparation for mass spectrometry, leaving only incompletely cleaved peptides amenable for detection. This may explain why prior mass spectrometric analyses have not detected this protein in tissues with relatively low frequency of RPL39L-expressing cells. If RPL39 and RPL39L carry their functions as part of ribosomes, as has been suggested before ^4, 39^, the RPL39L/RPL39 ratios that we estimated imply that ∼1% of the ribosomes in pluripotent cells contain RPL39L. Moreover, single cells that contain *RPL39L* reads also contain *RPL39* reads, which indicates that pluripotent cells have heterogeneous RPL39/RPL39L ribosome populations. The relatively high expression of *RPL39L* mRNA in extravillous trophoblast, breast glandular and pancreatic endocrine cells from the Human Protein Atlas, as well as multiple cancer types, suggested that RPL39L may support protein synthesis in the context of secretory processes.

By morphological and molecular characterization of KO cell lines, we demonstrated that RPL39L contributes to mESC pluripotency and differentiation along multiple lineages. Like RPL10L, RPL39L is a recently evolved autosomal chromosome-encoded paralog of an X chromosome-encoded RP. These paralogs are thought to take part in the formation of testis-specific ribosomes, necessary for spermatogenesis ^40^. Indeed, RPL10L has been shown to compensate for the reduced level of RPL10 during meiotic sex chromosome inactivation ^3^, while mice deficient in RPL39L exhibit spermatogenesis defects ^4, 11^. We found that *RPL39L* KO mESCs are not only impaired in the ability to differentiate into sperm cells but also show defects in spontaneous differentiation. The expression of ectoderm markers was increased and of endoderm markers decreased in embryoid bodies derived from *RPL39L* KO mESCs compared to those derived from WT cells. Whether these defects that we observed *in vitro* can also be detected in the early mouse development has not been investigated ^4, 11^.

To evaluate the impact of RPL39L on translation, we carried out ribosome profiling ^16^. Consistent with previous studies ^4, 11^, we found a small upregulation in the polysome-to-monosome ratio in the KO compared to WT clones, which indicates an increased protein synthesis rate. At the level of individual mRNAs, the changes in translation efficiency were very consistent across KO clones and occurred in mRNAs with specific functions. ER and Golgi-associated membrane proteins were generally less translated, while stress response and oxidative damage-related proteins were more translated in the *RPL39L* KO compared to WT mESCs. The increased phosphorylation of PERK and EIF2A further indicated that the stress was due to the ER-associated unfolded protein response (UPR) ^41^, consistent with an apparent increase in protein degradation that we also consistently found across all *RPL39L* KO clones. Pluripotent cells have higher resistance to ER stress than differentiated cells. However, the normal differentiation along the endodermal and ectodermal lineages requires UPR to be maintained at a specific level ^42^, which may explain why differentiation is perturbed as the mESCs experienced increased ER stress upon *RPL39L* KO.

We demonstrated that the reduced protein levels in *RPL39L* KO relative to WT mESCs were restored by the inhibition of proteasome and autophagy, showing that the protein stability is impaired in *RPL39L* KO cells. Some of the destabilized proteins have a testis bias of expression (according to the mRNA level data in the HPA), but the set more generally included proteins that are involved in cytoskeleton organization, cellular localization (specifically to membranes) and polarization. As we did not find an increased frequency of amino acid misincorporation in the KO clones, we investigated whether the decreased protein stability is due to alterations in other co-translational processes such as folding. RPL39 is known to interact with transmembrane alpha helices that fold in the NPET, but not with the alpha helices of secretory proteins ^43^. To elucidate the origin of these interactions, we turned to structural analyses. Obtaining pure populations of RPL39L ribosomes is very challenging and has not been achieved so far. Thus, to determine how RPL39/RPL39L impact the NPET structure we overexpressed the mouse RPL39 and RPL39L in *RPL39* KO yeast strains. The *RPL39* KO yeast lines are sensitive to cold, paromomycin and AZT, phenotypes that have been described before ^21, 22^. These phenotypes are rescued by both mouse RPL39 and mouse RPL39L, which indicates that RPL39L can perform the basic functions of RPL39. Analysis of single ribosome particles from these RPL39/RPL39L-expressing strains revealed that the main structural difference is a conformational change that involves the conserved residue I35, and leads to the appearance of a hydrophobic patch in the vestibular region of NPET, in the position where RPL39-containing ribosomes expose a positive charge. The loop containing Q28 and M36 further shows a higher flexibility compared to that provided by RPL39. Notably, we did not find electron densities supporting an occlusion of the tunnel region by RPL39L as proposed by a previous study ^4^. In fact, we did not find such evidence upon re-examining the published data either. Thus, it seems unlikely that RPL39L promotes the folding of specific proteins by occluding the NPET. What other mechanism could be responsible? Early studies hypothesized that a hydrophobic patch near the vestibular region of the NPET nucleates the co-translational folding of alpha helices ^8^. While the first structures of the 60S subunit of an archeal ribosome ^44^ did not identify such a patch ^45^, in prokaryotes, the uL23 RP, located at the vestibular region of the NPET recruits the trigger factor a (TFa) ^46^ to provide the hydrophobic surface for folding of cytosolic hydrophobic proteins ^47^. In eukaryotes, RPL39 replaces a loop of uL23 in the lower part of the tunnel, making it considerably narrower ^48^, which is thought to promote the entropic stabilization of alpha helices ^49^, and probably reducing the need for the hydrophobic patch to promote nascent protein folding. However, a narrower tunnel also leads to a reduced speed and increased accuracy of translation, both of which are impacted by the deletion of RPL39 ^21^. It is remarkable that the distinguishing feature of RPL39L ribosomes is the re-emergence of the hydrophobic patch, provided by RPL39L instead of the TFa chaperone, as chaperone function has diversified during the evolution of eukaryotes ^50^.

Although our study was done in mESCs, we found that the RPL39L-sensitive proteins have a testis bias of expression. To further understand the properties of these RPL39L-sensitive proteins, we further analyzed the dynamics of translation by calculating ribosome dwell time on individual codons and on codons encoding individual amino acids. We found that in *RPL39L* KO cells the dwell time was increased on leucine codons and reduced on histidine codons. These amino acids occur in helix-turn-helix, zinc finger, and leucine zipper coiled coil motifs ^51^, which are found in amphiphilic alpha helices ^52, 53^. The mRNA level data from HPA further shows that proteins with coiled-coil domains that are destabilized in the *RPL39L* KO mESCs exhibit an expression pattern surprisingly similar to that of *RPL39L,* in that they have highest mRNA-level expression in the spermatogenic lineage and/or trophoblast. Although amphiphilic alpha helices can fold in RPL39-containing ribosomes, the presence of a hydrophobic patch and better hydrophilic/hydrophobic partitioning may allow a more reliable folding of such domains, important when the burden of protein production is high, as in secretory cells (glandular, endocrine etc.) or cells that undergo large changes in polarity (developing sperm cells, ES cells etc) ^54, 55^.

Our work elucidates the function of a specialized ribosome, defined by RPL39L and present in sperm cells, but also other cell types such as mESCs. However, additional questions remain to be addressed in the future. First, the implications of *RPL39L* KO on the *in vivo* animal development are insufficiently understood. As already mentioned, RPL39L-deficient mice obtained via CRISPR/Cas9 genome editing ^4, 11^ were not investigated beyond the spermatogenesis defect. At the molecular level, it will be interesting to investigate further how ES cells and cancer cells fine-tune their translation output and the quality of synthesized proteins using modular ribosome components.

## Acknowledgements

We are grateful to Dr. Constance Ciaudo for suggestions regarding mESC differentiation, to Dr. Arnaud Schieberich for the hBM-MSC cells, Dr. Frederic Schmitt for the mouse samples, Dr. Sebastian Hiller and Dr. Nenad Ban for insightful discussions on the ribosome structure, and Dr. Vinay Tergaonkar for suggestions on the project. We thank the Imaging Core Facility at Biozentrum (IMCF), especially Laurent Guerard, Dr. Sara Roig and Dr. Kai Schleicher for help with the image acquisition and data analysis, and the sciCORE team for supporting the computational infrastructure. We would also like to thank Stella Stefanova and Janine Boegli of Biozentrum FACS core facility for help with the FACS data acquisition, and Philippe Demougin and Dr. Christian Beisel from the genomics facility Basel for help with RNA sequencing. Cryo-EM data were collected in the Scientific Center for Optical and Electron Microscopy (ScopeM). We thank Miroslav Peterek (ScopeM) for support and Pavel Afanasyev (CEMK) for helpful discussions. This work was supported in part by the SNF grant #51NF40_141735 for the NCCR project “RNA & Disease’’ in which the Zavolan group participated. The results shown here are in part based upon data generated by the TCGA Research Network: https://www.cancer.gov/tcga.

## Methods

### Quantification of RP gene expression in public RNA-seq data

Raw gene expression counts generated by STAR ^56^ for all 11,274 samples of the TCGA projects were downloaded from the GDC portal (https://portal.gdc.cancer.gov/). In addition, STAR 2-PASS genomic alignments (BAM format) of short reads from 1226 breast cancer and solid normal tissue samples of the TCGA-BRCA project were obtained from the GDC portal (accession number phs000178.v11.p8). Short reads from a single cells study of embryogenesis ^10^ were downloaded in .fastq format from the NCBI Sequence Read Archive (SRA, https://www.ncbi.nlm.nih.gov/sra) and aligned with STAR v2.7.10b in 2-PASS mode. Normalized gene TPM values across cell types and tissues estimated from pseudo-bulk scRNA-seq data (https://www.proteinatlas.org/download/rna_single_cell_type_tissue.tsv.zip) and cell-level gene count data (https://www.proteinatlas.org/download/rna_single_cell_read_count.zip) were downloaded from the HPA portal ^9^.

From 90 RPs expressed in human cells, a subset of 4 core RPs (RPS9, RPS14, RPL4, RPL32) was stringently selected based to satisfy the following criteria: (1) are deeply embedded into the rRNA ^57^, (2) bind early to ribosomal subunits during synthesis and assembly ^58^, (3) are present in other taxonomic groups apart from eukaryotes ^59^, (4) do not have strong paralogs in the human genome with DIOPT score>2 and “high” DIOPT rank ^60^, (5) do not display strong evidence of tissue-specific expression ^2^. RPs with these features are more likely to be essential for cell survival (Fig. S1A), as indicated by significantly lower scores in CRISPR ^61^ and RNAi screening ^62^ (data available at the DepMap portal https://depmap.org/portal/).

mRNAs encoding RP genes were extracted from the v36 version of the GENCODE comprehensive annotation of the hg38 human genome assembly ^63^. RNA-SeQC ^64^ was utilized to obtain raw gene counts from uniquely mapped reads (MAPQ=255) as well as from all reads, including multi-mapped ones (MAPQ>=0), in bulk RNA-seq data from the TCGA-BRCA project and scRNA-seq data from the embryogenesis study ^10^. Gene counts of *RPL39L* and of core RPs comprised only a small fraction (<7%) of multi-mapped reads in breast cancer and normal tissue samples (Fig. S1B), consistent with these RPs having a relatively low number of processed pseudogenes ^65^. Gene lengths of the RPs were estimated as average transcript lengths weighted by TPM values. StringTie v.2.2.1 ^66^ was utilized to identify mRNAs and quantify their TPM values across TCGA-BRCA samples. Median gene lengths of *RPL39L*, *RPL39*, and core RPs were checked and found to have only minor differences between cancer and normal samples, indicating minimal isoform variation (Fig. S1C). The arithmetic means of these median values were further used as gene length estimates.

For *RPL39L, RPL39* and core RPs, reads-per-kilobase (RPK) values were estimated from public bulk RNA-seq data from the TCGA project and scRNA-seq data from the embryogenesis study ^10^ as raw gene counts with pseudocount=1 divided by the gene length. RPL39-to-core RP and RPL39L-to-core RP ratios were calculated as the ratios of RPL39 (respectively, RPL39L) RPK values relative to the median of core RP RPK values.

### Cell culture

In the absence of feeder cells, the WT E14 mESC line was cultured in Dulbecco’s Modified Eagle Media (DMEM) (Gibco 31966), which contained 20% fetal bovine serum (FBS; Gibco 16141079) tested for optimal mESC growth, NEAA (Gibco 11140050), sodium pyruvate (Gibco 11360039), 100 U/mL LIF (Millipore ESG1107) and 0.1 mM 2-ß-mercaptoethanol (Merck ES-007-E), on 0.2% gelatin-coated plates. The culture medium was changed daily, and the cells were passaged every second or third day. Cells were cultured at 37°C in 5% CO2.

### RPL39L CRISPR in E14.tg2a (E14) mouse embryonic stem cells

All transfections for the CRISPR knockout were performed using the lipofectamine 2000 reagent (Life technologies), according to manufacturer’s instructions. SgRNAa for the *RPL39L* gene were designed in pairs from the sequences upstream and downstream of the 5’ and 3’UTRs, respectively. Upstream sgRNAs were cloned into the px330 plasmid backbone with an mCherry marker, and downstream sgRNAs were cloned in the px330 backbone with a GFP marker. The cells positive for both mCherry and GFP expression were FACS-sorted. For single cell colony selection, FACS sorted cells were diluted to a concentration of 0.75 cells per 100 μl in the ESC culture media. 100μl of this solution was added to each well of a 96 well plate. The cells were allowed to grow for 2 days before wells with single clones were marked. Forward primers for CRISPR KO validation were designed upstream of the upstream sgRNAs and the reverse primers were designed downstream of the downstream sgRNAs (see Table S1). In clones with homozygous KO the PCR product should consist of a single band, migrating lower than the wild type band (692bp and 587bp in KO, compared to 1157bp in WT) when run on a 1.3% agarose gel. The bands were excised and gel purified using the Qiagen Gel purification Kit, according to manufacturer’s instructions. The purified DNA was cloned into a pUC19 plasmid backbone using Zero Blunt Topo kit (ThermoFischer) according to manufacturer’s instructions. This plasmid was sequenced to validate the complete excision of *RPL39L*.

### LC-MS analysis

#### Sample preparation

Cells were collected and lysed in 50 μl lysis buffer (2 M Guanidinium-HCl, 0.1 M HEPES, 5 mM TCEP, pH = 8.3) using strong ultra-sonication (10 cycles, Bioruptor, Diagnode). Protein concentration was determined by BCA assay (Thermo Fisher Scientific 23227) using a small sample aliquot. Sample aliquots containing 50 μg of total proteins were supplemented with lysis buffer to 50 µl, reduced for 10 min at 95 °C and alkylated at 10 mM iodoacetamide for 30 min at 25 °C followed by adding N-acetyl-cysteine to a final concentration of 12.5 mM to quench any iodoacetamide excess. For global proteomics analyses, proteins were directly digested by incubation with sequencing-grade modified trypsin (1/50, w/w; Promega, Madison, Wisconsin) overnight at 37°C. For targeted analysis of RPL39 and RPL39L, a mixture containing 100 fmol of heavy reference peptides were added to the samples. Then, protein samples were propionylated by adding N-(Propionyloxy)-succinimide (0.15 M in DMSO) to a final concentration of 22.5 mM and incubating for 2h at 25°C with shaking at 500 rpm. This step prevents trypsin from cleaving after lysine residues and leads to the generation of larger peptides suited for LC-MS analysis for our target proteins. To quench the labeling reaction, 1.5 μL aqueous 1.5 M hydroxylamine solution was added and samples were incubated for another 10 min at 25°C shaking at 500 rpm. Subsequently, the pH of the samples was increased to 11.9 by adding 1 M potassium phosphate buffer (pH 12) and incubated for 20 min at 25°C shaking at 500 rpm to remove propionic acid linked to peptide hydroxyl groups. The reaction was stopped by adding 2 M hydrochloric acid until a pH < 2 was reached. After adding 70 ul of a 1 M TEAB (pH=8.5), proteins were digested by incubation with sequencing-grade modified trypsin (1/50, w/w; Promega, Madison, Wisconsin) overnight at 37°C. For all samples, the generated peptides were cleaned up using iST cartridges (PreOmics, Munich, Germany) according to the manufacturer’s instructions. Samples were dried under vacuum and stored at −80 °C until further use.

#### Global proteomics LC-MS analysis

Dried peptides were resuspended in 0.1% aqueous formic acid and subjected to LC–MS/MS analysis using a Q Exactive HF Mass Spectrometer fitted with an EASY-nLC 1000 (both Thermo Fisher Scientific) and a custom-made column heater set to 60°C. Peptides were resolved using a RP-HPLC column (75μm × 30cm) packed in-house with C18 resin (ReproSil-Pur C18–AQ, 1.9 μm resin; Dr. Maisch GmbH) at a flow rate of 0.2 μLmin-1. The following gradient was used for peptide separation: from 5% B to 15% B over 10 min to 30% B over 60 min to 45 % B over 20 min to 95% B over 2 min followed by 18 min at 95% B. Buffer A was 0.1% formic acid in water and buffer B was 80% acetonitrile, 0.1% formic acid in water.

The mass spectrometer was operated in DDA mode with a total cycle time of approximately 1 s. Each MS1 scan was followed by high-collision-dissociation (HCD) of the 10 most abundant precursor ions with dynamic exclusion set to 30 seconds. For MS1, 3e6 ions were accumulated in the Orbitrap over a maximum time of 100 ms and scanned at a resolution of 120,000 FWHM (at 200 m/z). MS2 scans were acquired at a target setting of 1e5 ions, maximum accumulation time of 100 ms and a resolution of 30,000 FWHM (at 200 m/z). Singly charged ions and ions with unassigned charge state were excluded from triggering MS2 events. The normalized collision energy was set to 35%, the mass isolation window was set to 1.1 m/z and one microscan was acquired for each spectrum.

#### Targeted LC-MS analysis of RPL39L

Parallel reaction-monitoring (PRM) assays ^67, 68^ were generated from a mixture containing 25 fmol/µL of each proteotypic N-(Propionyloxy)-succinimid labeled heavy reference peptide (SSHKTFR (RPL39 human and mouse), SSHKTFTIKR (RPL39L human), ASHKTFR (rpl39l mouse), JPT Peptide Technologies GmbH). 2 µL of this standard peptide mix were subjected to LC–MS/MS analysis using a Q Exactive plus Mass Spectrometer fitted with an EASY-nLC 1000 (both Thermo Fisher Scientific) and a custom-made column heater set to 60°C. Peptides were resolved using a EasySpray RP-HPLC column (75μm × 25cm, Thermo Fisher Scientific) and a pre-column setup at a flow rate of 0.2 μL/min. The mass spectrometer was operated in DDA mode. Each MS1 scan was followed by high-collision-dissociation (HCD) of the precursor masses of the imported isolation list and the 20 most abundant precursor ions with dynamic exclusion for 20 seconds. Total cycle time was approximately 1 s. For MS1, 3e6 ions were accumulated in the Orbitrap cell over a maximum time of 50 ms and scanned at a resolution of 70,000 FWHM (at 200 m/z). MS1 triggered MS2 scans were acquired at a target setting of 1e5 ions, a resolution of 17,500 FWHM (at 200 m/z) and a mass isolation window of 1.4 Th. Singly charged ions and ions with unassigned charge state were excluded from triggering MS2 events. The normalized collision energy was set to 27% and one microscan was acquired for each spectrum.

The acquired raw-files were searched using the MaxQuant software (Version 1.6.2.3) against the same human and mouse database mentioned above using default parameters except protein, peptide and site FDR were set to 1 and Lys8, Arg10 and propyl (K) were added as variable modifications. The search results were imported into Skyline (v21.1.0.278)^69^ to build a spectral library and assign the most intense transitions to each peptide. An unscheduled mass isolation list containing all peptide ion masses was exported and imported into the Q Exactive Plus operating software for PRM analysis. For PRM-MS analysis, peptide samples were resuspended in 0.1% aqueous formic acid. Due to the required protein propionylation for rpl39 and rpl39l LC-MS analysis, the heavy reference peptides were already spiked in at a concentration of 2 fmol of heavy reference peptides per 1 µg of total endogenous peptide mass during sample preparation (see above). The samples were subjected to LC–MS/MS analysis on the same LC-MS system described above using the following settings: The MS2 resolution of the orbitrap was set to 17,500/140,000 FWHM (at 200 m/z) and the fill time to 50/500ms for heavy/light peptides. AGC target was set to 3e6, the normalized collision energy was set to 27%, ion isolation window was set to 0.4 m/z and the first mass was fixed to 100 m/z. A MS1 scan at 35,000 resolution (FWHM at 200 m/z), AGC target 3e6 and fill time of 50 ms was included in each MS cycle. All raw-files were imported into Skyline for protein / peptide quantification. To control for sample amount variations during sample preparation, the total ion chromatogram (only comprising precursor ions with two to five charges) of each sample was determined using Progenesis QI software (Nonlinear Dynamics (Waters), Version 2.0) and used for normalization of light (endogenous) peptide abundances.

### Sample preparation for inhibition of protein degradation

E14 and RPL39L KO cells were treated for 5h at 37°C and 5% CO2 with 10μM (S)-MG132 (STEMCELL Technologies Catalog #73264) and 5μM Bafilomycin-A1 (STEMCELL Technologies Catalog #74242) in the normal media described above. For LC-MS as well as for western blot analysis, cells were scraped and washed twice with warm DPBS, then lysed in a corresponding lysis buffer.

### Spontaneous differentiation

Spontaneous differentiation of mESC lines was carried out in conventional mESC medium (as stated above) with 10% fetal bovine serum and no LIF ^70^. To avoid attachment, embryoid bodies (EBs) were generated by growing 750’000 cells in suspension for 6 days in non-adherent dishes (Greiner Bio-One 633181). Spheroids were allowed to grow without disturbing the plate for the first 3 days. The medium was changed every second day afterwards. The EBs were harvested after 6 days with 25ml Pasteur pipettes and washed with 1X PBS thrice before further analysis.

### Spermatogenic differentiation

Differentiation of mESC toward sperm cells was done according to the protocol described in ref. ^12^. In short, E14 cells were harvested and plated as a hanging media drop on Petri dishes filled with PBS on the bottom of the plate, with 1250 ESC per 25 μl in each drop. The resultant EBs were transferred onto Petri dishes (10–15 EB per dish) after 3 days in hanging drop culture. Neurobasal medium (Gibco 21103049), supplemented with B27 (Invitrogen 17504044) was used as the differentiation medium, according to the manufacturer’s protocol 0.1 µM retinoic acid final concentration (Sigma R2625-50MG) was added to the culture medium one day after EB transfer, and the medium was replaced every two days to minimize deterioration. The contents of each plate were harvested after 4 days.

### Polysome profiling and ribo-seq

For ribo-seq analysis, WT and *RPL39L* KO E14 mESCs were propagated in 5×15cm Petri Dishes (Falcon, 353025) per sample as described in the cell culture section. The medium was replenished three hours prior to cell collection at the confluency of 50-70%. Before harvesting, the cells were treated with 100μg/ml cycloheximide (CHX) (Sigma, G7698) for 15 minutes at 5% CO2, 37°C, to freeze the elongating ribosomes. The cells were harvested on ice in a cold room. Cells were washed twice with ice cold DPBS (Lonza, BE17-512Q) containing 100μg/ml CHX. Cells were scraped, collected, spun down, flash frozen and stored at −80 °C.

The cell pellet was resuspended in 900μl of ice-cold polysome lysis buffer (20 mM Tris-HCl (Sigma, T294), pH=7.5; 100 mM NaCl (Sigma, 71386); 10 mM MgCl2 (Sigma, 63069); 1% Triton X100 (Sigma, T8787); freshly added 2 mM DTT (Sigma, 646563); 100 µg/ml cycloheximide (Sigma, G7698); 400U of RNAsin plus RNase inhibitor (Promega, N261B); 20U of Turbo DNase (Ambion, AM2238) and Complet mini, EDTA-free protease inhibitors (Roche, 11836170001)) by pipetting up and down. After resuspending the pellet, the sample was incubated for 5 minutes at 4°C with continuous rotation (50 rpm), passed through a 23G needle (Braun, 4657640) for minimum 10 times, followed by additional 5 min incubation at 4°C with continuous rotation (50 rpm). The cell lysate was clarified by centrifugation at 3000 g/3min/4°C followed by centrifugation of supernatant from the first step at 10’000g/5min/4°C. 50 µl of clarified lysate from each sample was kept aside for mRNAseq library preparation and was snap frozen and stored at −80 °C. The optical density (OD) of remaining lysate was measured at A_260_ with a NanoDrop2000. For ribo-seq analysis, the lysate equivalent to OD (A_260_) = 7 was subjected to RNase I digestion ( 5U per OD, 35U in total; Invitrogen, AM2294) at 22 °C/ 20 minutes with continuous rotation at 1000 rpm in thermoblock (Eppendorf). RNase I was inactivated by adding 10µl of RNase inhibitor SuperaseIN (Invitrogen, AM2696) in each reaction. An equal amount of undigested lysate was also used for the polysome profile.

10-50% linear sucrose gradient was prepared using a Gradient Master instrument (Biocomp) according to the manufacturer’s instructions. In brief, 10 and 50% sucrose (Sigma, 84100) solution was prepared in buffer containing 50 mM Tris-HCl, pH 7.5 (Sigma, T294); 50 mM NH4Cl (Sigma, 09718); 12 mM MgCl2 (Sigma, 63069); 100 µg/ml CHX (Sigma, G7698); 0.5mM DTT (Sigma, 646563) and 10 µL SuperaseIN (Invitrogen, AM2696). A 14×89mm tube (Beckman Coulter, 331372) (used in rotor SW-41/TH-641) (Beckmann Coulter) with a long cap that can hold 800 µl of sample was used to prepare the gradient. First 10% solution was laid in the tube followed by 50% solution that was under-laid with the help of a long syringe. The gradient was prepared by using a pre-program for 10% to 50% sucrose gradient in Gradient master 108 instrument (Biocomp). The gradient was cooled down at 4°C by keeping it in the fridge for a minimum of one hour. Undigested (for polysome profile) and digested samples (for ribo-seq) were loaded onto pre-cooled 10-50% sucrose gradient and centrifuged at 35,000 rpm for 3 hrs at 4°C in a SW-41Ti rotor (Beckmann Coulter). Finally, all gradients were monitored at wavelength A_254_ and fractionated using Piston Gradient Fractionator (Biocomp). 30 separate fractions of 0.37 ml were collected in 1.5 ml Eppendorf tubes for each digested and undigested ribosome profiles using a Gilson collector attached with the fractionator.

The appropriate fractions containing 80S monosomes were processed for ribo-seq library preparation by combining the protocol from ^16, 71^. In brief, RNA was isolated from the appropriate monosomes fraction by using the hot phenol method. RNA fragments of appropriate size (28-32 nt) were obtained by running samples on 15% polyacrylamide denaturing TBE-Urea gel and visualized by SYBR Gold dye (Life Technologies). Size selected RNA was dephosphorylated by T4 polynucleotide kinase (PNK, New England Biolabs, B0201S) treatment for 1 hour at 37°C. PNK was heat inactivated and RNA was purified using phenol chloroform method and overnight precipitation of RNA in ethanol. An RNA amount equivalent to 12 ng was used to prepare the sequencing library using SMARTer® smRNA-Seq Kit for Illumina® (Takara 635031) according to the kit manual till cDNA synthesis. After cDNA synthesis, rRNA was depleted by using the protocol and probe defined in ref. ^16^, followed by final PCR according to Smarter kit with multiplexing barcodes. The PCR product was purified on 8% polyacrylamide native TBE gel and sequenced on a NextSeq 500 instrument at genomic facility Basel.

The translation rate of cells was calculated from polysome profiles, as the area under the curve corresponding to polysomes divided by the area under the curve corresponding to the monosome (80S). This ratio was calculated in every sample/clone and compared to the WT control. The unpaired one-sided t-test was used to determine whether the KO clones exhibit significantly increased rate of translation relative to WT.

### RNA-seq sample preparation

Cells were maintained as described in the cell culture section. RNA was isolated using AccuPure Cell/Blood RNA Mini Kit (96) (AccuBioMed, R10096) using a iColumn24 Robot (AccuBioMed) with DNase1 treatment and 50μl volume elution. RNA-seq samples were prepared using the Trueseq Standard mRNA Illumina kit and sequenced on a NovaSeq 6000 instrument at the genomics facility Basel.

### Analysis of ribosome profiling data

Reads from fastq files were trimmed with fastx_clipper from FASTX-Toolkit version 0.0.14 with parameters ‘-a (3’adapter) AAAAAAAAAA, -l (minimum-length) 20, -c (discards non-clipped sequences) and -n (discards sequences with unknown (N) nucleotides)’. The trimmed reads were further trimmed with fastq_quality_trimmer from the same toolkit with -t (minimum quality) 20, -Q (quality type) 33. Then the trimmed reads were filtered with fastq_quality_filter from the same toolkit for read quality with the following parameters: ‘-q (minimum quality) 20, -p (minimum percent of bases that must have [-q] quality) 90’, -l (minimum-length) 20. These reads were first aligned to ribosomal RNA (rRNA) sequences obtained from *Mus musculus* ribosomal DNA (rDNA), complete repeating unit (https://www.ncbi.nlm.nih.gov/nuccore/bk000964) using Segemehl ^72^ version 0.2.0. The reads that did not map to rDNA were then aligned to the longest coding transcripts for each gene identified from *Mus musculus* GRCm38–mm10 genome assembly, Ensembl 99 annotation using Segemehl. The uniquely mapped reads from this alignment were used for downstream analysis.

### Analysis of RNA sequencing data

Single-end reads from raw fastq files were processed using ZARP ^73^ workflow with the default parameters, *Mus musculus* GRCm38–mm10 genome assembly, Ensembl 99 annotation, 3’ adapter ‘GATCGGAAGAGCACAC’, ‘SR’ for library type (strand-specific reads coming from the reverse strand). The kallisto ^74^ output (version 0.46.2) of ZARP workflow was used for downstream analysis.

### Analysis of differential expression, translation and translational efficiency

Differential expression (RNA-seq) and differential translation (ribo-seq) analyses were performed using the Deseq2 R package ^75^ version 1.34.0 with default parameters. The deltaTE ^76^ procedure was applied for differential translation efficiency analysis using the reads mapped to coding sequence (CDS) regions obtained from GRCm38–mm10 genome assembly, Ensembl 99 annotation both for RNA-seq and Ribo-seq libraries. Rsubread ^77^ R package version 2.8.2 was employed to obtain the RNA-seq reads aligned to CDS regions. An *in-house* algorithm was used to obtain the ribo-seq reads mapped to CDS regions based on their estimated P-sites.

### Gene Ontology (GO) analysis

The ClusterProfiler ^78^ R package version 3.18.1 was used for all the GO term analyses reported in this study. ComplexHeatmap ^78, 79^ version 2.6.2 and circlize ^80^ version 0.4.15 R packages were used to construct the heatmaps. InteractiVenn ^81^ was used for Venn diagrams.

### Analysis of LC-MS data

The raw files were searched against a protein database containing sequences of the SwissProt ^82^ entries of *Mus musculus* (in total 17,137 protein sequences) along with commonly observed contaminants using FragPipe version 18.0.0 (downloaded from https://github.com/Nesvilab/FragPipe/releases). Protein intensities obtained from the FragPipe platform were imputed with the impute.knn function from *impute* R package ^83^ version 1.64.0. Normalization of protein intensities and differential protein expression analyses were performed using *limma* R package ^84^ version 3.64.0.

For amino acid mismatch analysis, the procedure developed by Mordret et al.^19^ was employed. The dependent peptides required for mismatch identification were obtained using MaxQuant computational platform ^85^ version 2.1.3.0.

### RT-PCR

RNA from Spontaneous and Spermatogenic differentiation was isolated using AccuPure Cell/Blood RNA Mini Kit (96) (AccuBioMed, R10096) using iColumn24 Robot (AccuBioMed) with DNase1 treatment and 50ul volume elution. Reverse transcription was performed from 500-1000ng of RNA using SuperScript IV First-Strand Synthesis System (Invitrogen, 18091200) using Random hexamers (Promega, 300453). qPCR reaction was performed in 20ul volume using PowerSYBR Green PCR Master Mix (AppliedBiosystems, 4367659) and QuantStudio 3 System (Applied Biosystems, A28567) using Comparative CT (ΔΔCT) analysis and Standard Run mode. Rrm2 was treated as endogenous control for all analyses. The primer sequences used are in Table 2.

### Spontaneous differentiation organoids immunofluorescence

Organoids were fixed using 4% PFA in PBS at 4 °C for 2hrs. After two washes in PBS, the organoids were transferred to 10% sucrose solution in PBS and stored at 4 °C for 1 day. This was followed by a 1-day incubation in 20% sucrose in PBS at 4 °C, and finally by a 1-day incubation in 30% sucrose in PBS at 4 °C. Next, the organoids were embedded in PolyFreeze Tissue Freezing Medium (Sigma, SHH0025) in ibidi slides (ibiTreat, 80826), snap frozen on dry ice and stored at −80 °C until cryosectioning. Cryosections of 12-μm thickness were made on Superfrost Ultra Plus Gold Adhesion slides (Thermo Fisher, 11976299) using a Leica Microsystems cryostat. Slides were stored at −80 °C or processed immediately. For IF, slides were air-dried at RT for 1 h, washed in PBS three times 5 min each and permeabilized in 0.2% Triton X-100 in PBS for 30 min at RT. The cryosections on each slide were then circumscribed using ImmEdge Hydrophobic Barrier Pen (Vector Labs) and blocking solution (1% BSA, 5% mouse serum in 0.2% Triton X-100) was added for 30 min at RT. Primary antibodies against Gata4-Alexa594 (SantaCruz, sc-25310) and Nestin-Alexa488 (SantaCruz, sc-23927) were diluted 1:50 in blocking solution (with 0.1% Tween) and incubated in dark at 4 °C overnight. In the end slides were washed 3 times with 0.1%Tween in PBS and mounted with VECTASHIELD Antifade Mounting Medium with DAPI (Vector Laboratories, H-1200-10). The images were acquired using Zeiss LSM800 confocal microscope and analyses using Fiji software ^86^ and visualized and deposited using OMERO ^87^ (project ID:12920).

The image analysis was done in Fiji ^86^ using software available on github https://github.com/imcf-shareables/stem_cell_analysis. The script uses TrackMate ^88^ and StarDist ^89^ to segment the nuclei in 3D in a defined region of interest. The mean intensity of the endoderm marker was measured in the nuclear region while the mean intensity of the ectoderm marker was measured in a 3D layer around the nuclei obtained by dilating the nuclei and subtracting the originals from the dilation, using the 3DImageJSuite ^90^. The results were then saved to CSV for statistical analysis.

### Western blotting

Samples were collected from cell culture by scraping, centrifuged (5min @210rcf, 4℃), washed and resuspended in RIPA buffer containing Phosphatase inhibitor (Roche 4906837001) and protease inhibitor cocktail (Roche 118361530001). After sonication (2x 4 times Amp60%, pulse 0,5) on Hilsher UP50H probe sonicator and centrifugation (5min @ 5000rpm, 4℃) of the samples, a BCA measurement was executed to determine the protein concentration of the samples. For the SDS-Gel electrophoresis, 30ug of the samples were mixed with 4x Lämmli Buffer containing b-mercaptoethanol, heated up for 10 minutes at 95°C and centrifuged before loading onto a 10 well 5-20% gradient gel (BioRad 456-1093). The finished gels were blotted via semi dry method (20V, 400mA, 45min) onto a nitrocellulose membrane (GE, Amersham Protran Premium 0,2um NC), dyed with ponceau, washed with 1x TBST, blocked with 5% BSA in 1x TBST and kept overnight at 4°C rotating in the respective primary Antibody. The membranes were washed afterwards 3x 10min with 1xTBST then put into secondary Antibody at RT for 1h and washed again. They were imaged in the Fusion FX from VILBER (Software FUSIONFX7 Edge 18.11) with Amersham ECL Western Blotting Detection Reagents RPN2106 from GE Life science. For loading control the membranes were put in GAPDH (Histone H3 for EIF2A and P-EIF2A). For details of the antibodies and the respective concentrations used, please refer to Table S2.

### Yeast strains and media

Yeast strains were either grown in rich media composed of 1% w/v yeast extract, 1% (w/v) peptone, 40 mg/L-adenine, 2% (w/v) glucose (YPD) or in synthetic complete medium (HC) composed of 0.17% (w/v) yeast nitrogen base with ammonium sulfate and without amino acids, 2% (w/v) glucose and mixtures of amino acids (MP Biomedicals) depending on the auxotrophies used for selection. Cells were grown at 30°C or 23°C. Solid media contained 2% (w/v) agar and were supplemented with paromomycin (P9297 Sigma) or L-Azetidine-2-carboxylic acid (AZC; A0760 Sigma) when needed. Yeast RPL39 genomic deletion was done according to standard procedures [69], using pAG32 as PCR template for gene replacement and integration of the Hygromycin resistance gene confirmed by PCR (see Tables S3-S4).

### Yeast transformation

Three units of OD600 of yeast cells were grown in appropriate YPD or HC media to mid-log phase. Cells were spun down and washed in 1 volume of 1x TE and 10 mM LiAc. The pellet was then resuspended in 350 µL of transformation mix (1x TE, 100 mM LiAc, 8.5% (v/v) ssDNA, 70% (v/v) PEG3000), incubated with DNA (PCR product or 1µg of plasmid DNA) for 1 h at 42°C, spun down (30 sec at 10,000 xg at RT), resuspended in 100 μL of YPD or HC media and cells were plated onto selective media and incubated at 30°C.

### Plasmids

All RPL39 variants were cloned into the pRS413-GPD plasmid digested by the BamHI and SalI using the Gibson assembly kit (NEB). Guide blocks (IdT; Table S3) of 156 bp consisting of mouse RPL39, mouse RPL39L, or yeast mouse RPL39 and RPL39L were designed and used as PCR templates using dedicated primers (Table S4).

### Ribosome purification from yeast cells

The ribosomes from the various yeast lines were extracted using the protocol as described in ^91^. To summarize, the yeast cells were grown in HC -His medium and harvested by centrifugation at exponential growth phase and snap-frozen in liquid nitrogen. The frozen pellet was disrupted using SPEX SamplePrep 6875 Freezer Mill in liquid nitrogen. The crushed frozen pellet was resuspended in RES (50mM Hepes pH 7.6, 200 mM KCl, 10 mM MgCl2, 5 mM EDTA, 250 mM sucrose, 2 mM DTT). Centrifugation was used to clear cell debris for 60 minutes at a speed of 25.600 x g in a Beckmann Coulter Type 45 Ti rotor. The 80S ribosome-containing supernatant was decanted and added to a cushion of 50% (w/w) sucrose (62 mM Hepes pH 7.6, 62 mM KCl, 12 mM MgCl2, 6 mM EDTA, 50% (w/w) sucrose, 0.025% sodium azide, and 2 mM DTT), which was then centrifuged for 20 hours at 184.000xg and 4°C. (Beckman Ti70 rotor). The granules were resuspended in PRE buffer (50 mM Hepes pH 7.6, 10 mM KCl, 10 mM MgCl2, 0.02% sodium azide, and 2 mM DTT) after the supernatant was removed. Centrifugation at 103.000xg and 4°C for 14 hours separated the ribosomal subunits on a 50% (w/v) sucrose gradient (52 mM Hepes pH 7.6, 727 mM KCl, 10 mM MgCl2, 0.021% sodium azide, and 2 mM DTT) in Beckmann Coulter XE-90 ultracentrifuge.

### Cryo-EM analysis

#### Vitrification and cryo-EM data collection

Samples were vitrified on Quantifoil R2/2 holey carbon grids, coated in-house with 1 nm continuous carbon. Grids were subjected to glow discharge for 15 s with 15 mA current directly before sample vitrification. The climate chamber of a ThermoScientific Vitrobot was equilibrated to a temperature of 4°C and 95% humidity. A volume of 4 µl ribosome sample was subjected to a vitrification protocol using 30 s pre-blot incubation followed by 2 s or 3 s blotting, and subsequent rapid transfer into liquid ethane-propane mix. Micrographs were collected on a ThermoScientific Titan Krios cryo-electron microscope equipped with a Gatan K3 direct electron detector and a GIF BioContinuum energy filter. Images were acquired at 300 kV accelerating voltage in counted super-resolution mode with an electron dose of 45 e^-^/Å^2^ at a nominal magnification of 105000x, resulting in a pixel size of 0.84 Å/pixel. Micrographs were exposed for 1s and fractionated into 40 frames. The slit width of the energy filter was set to 20 eV.

#### Cryo-EM image processing

All image processing was carried out in cryoSPARC (version v3.3.1+220315) ^92^ (Fig. S7). Micrographs were dose weighted and motion corrected using Patch Motion Correction. Defocus was subsequently estimated by CTF fitting using Patch CTF Estimation. Particles were selected using the blob picker with a circular blob template with a diameter between 300 Å and 500 Å. Visual inspection confirmed that the blob picker accurately selected ribosomal particles. Particles were extracted with a box size of 600 pixels and scaled to 400 pixels (Nyquist limit 2.54 Å) and subjected to 2D classification into 200 classes. Classes that contained 60S or 80S ribosomal subunits were retained while empty classes and classes that contained junk particles (ice blobs, carbon edges), or 40S ribosomal subunits were removed. An initial model was generated from a subset of 63945 particles of dataset 1 (rpl39Δ-MmRPL39L) using cryoSPARC *ab-initio* model generation (1 class). A soft mask encompassing the 60S subunit, but excluding the 40S subunit was created from the initial model and used for refinement. The resulting map at 2.63 Å resolution and the associated mask were used as the initial model for refinement of all datasets. Using the reconstruction calculated from a subset of the data filtered to 30 Å resolution as the initial model, 3D maps of all samples were reconstructed by Homogeneous Refinement. As the resolution of the refined maps of samples Sc60S, rpl39Δ-ScRPL39, rpl39Δ-MmRPL39, and rpl39Δ-MmRPL39L exceeded the Nyquist limit of the binned particle images, particles were re-extracted with 600 pixel box size, without binning, and subsequently refined using Homogeneous Refinement and Non-uniform Refinement. Resolution of the maps was further improved by Global and subsequent Local CTF Refinement.

#### Model building and refinement

The yeast 60S ribosomal subunit structure (7TOO, ^93^) was docked into the density map by rigid-body fitting in UCSF Chimera ^94^. Parts of the original model that were outside of cryo-EM density due to flexibility were deleted from the model. Ribosomal protein and rRNA positions were adjusted in Coot ^95^ and MmRPL39 and MmRPL39L, respectively, were built into the density. Metal ions, chloride ions, water, spermine, and spermidine were placed into positive difference density according to the chemistry of the environment and the geometry of the coordinating groups; divalent ions were built as magnesium ions and monovalent ions were built as potassium ions. Metal ion coordination restraints were generated and optimized with ReadySet ^96^. The model was refined into the density using Phenix (dev-4788-00) ^97^ real space refinement (Fig. S7, Table S5), using five cycles with secondary structure, Ramachandran and side-chain rotamer restraints. The quality of the refinement was validated using real-space correlation coefficients (model vs. map at FSC=0.5). Images were prepared with PyMol (citation: The PyMOL Molecular Graphics System, Version 2.0 Schrödinger, LLC.).

## Data and code availability

Sequencing data has been deposited to the NCBI BioProject database, under accession PRJNA951511. Processing scripts are available from the zenodo repository, with DOI 10.5281/zenodo.7794007.

