## Supplementary Materials for "Ribosomal protein RPL39L is an efficiency factor in the cotranslational folding of proteins with alpha helical domains"

### Supplementary Figures

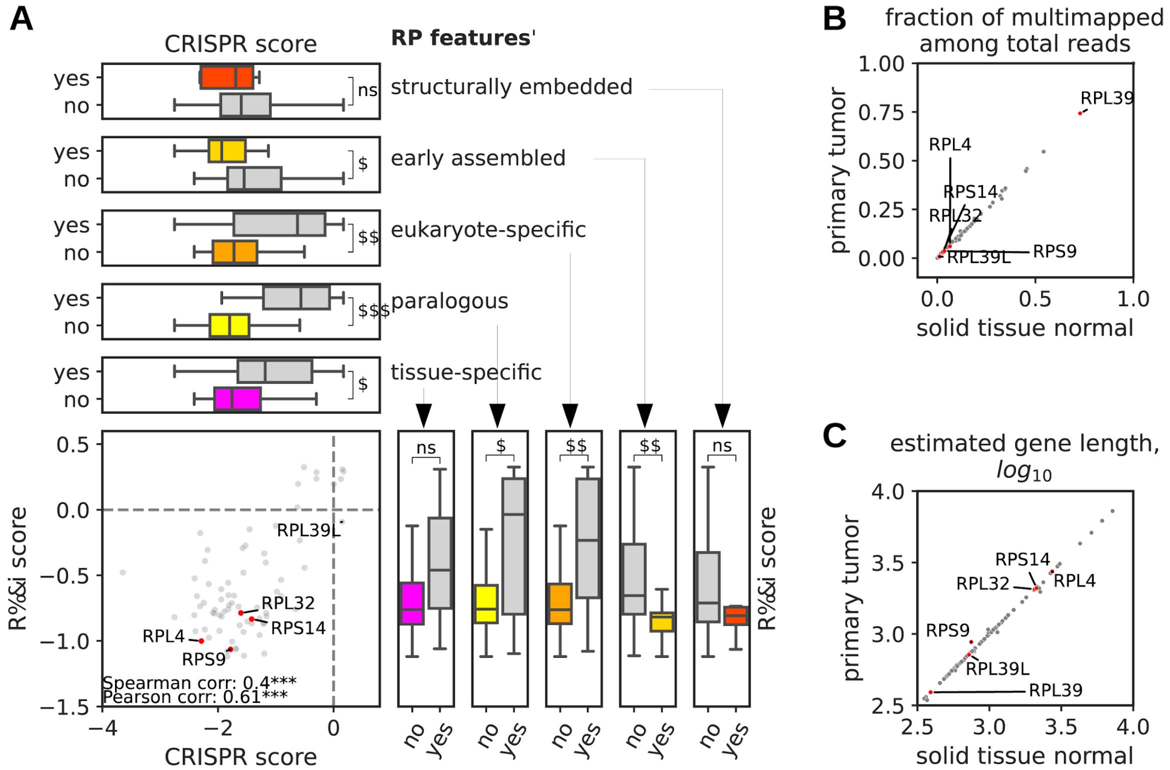


**Figure S1. Selection of core RPs. A.** Out of 90 RPs, 76 RPs with CRISPR and RNAi screening data available at the DepMap portal (https://depmap.org/portal/) were selected. For these, 90% quantiles of scores from individual cell line experiments (1078 CRISPR screens, 710 RNAi screens) are shown in a joint scatter plot (bottom left). The distributions of scores are compared between groups of RPs having or lacking each of the five features used to characterize the functional importance of RPs (see Methods, two-sided Mann-Whitney test, Bonferroni-adjusted for multiple testing). In all subpanels, ns, *, **, and *** correspond to p-values ≥0.05, <0.05, <0.01 and <0.001. **B.** Fraction of multi-mapped reads among all reads supporting the expression of each individual RP (see Methods). Median values of these fractions in normal tissue and primary tumor bulk RNA-seq samples of the TCGA-BRCA project are shown. RPL39 is one of the RPs with a very high fraction of multi-mappers. **C.** Estimated median gene length ($\log_{10}$) of expressed RPs in normal and primary tumor bulk RNA-seq samples of the TCGA-BRCA project.

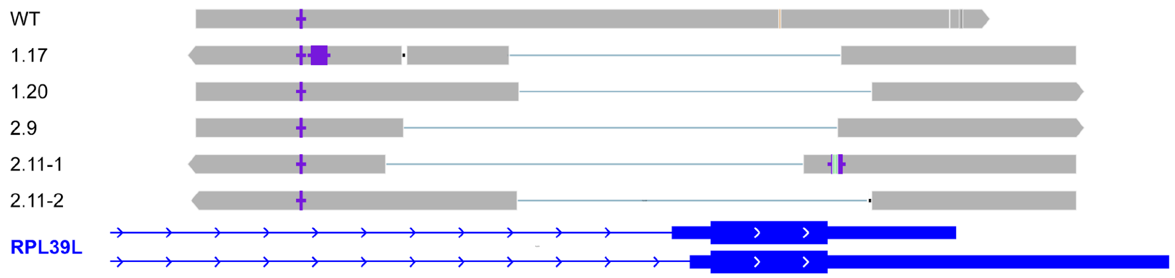


**Figure S2. Schematic representation of sequenced PCR amplification products derived from the *Rpl39l* locus of the *Rpl39l* KO lines.** PCR-amplified products of *Rpl39l* loci from extracted DNA of KO lines were sequenced. A single product was obtained from clones 1.17, 1.20 and 2.9, indicating homozygous deletions of *Rpl39l* coding region from the genome. In Clone 2.11, two products were obtained, one with a complete (2-11-2) the other with an almost complete (2-11-1) deletion of the coding region. The structure of the locus is shown in blue at the bottom on the panel, with two annotated transcripts that differ in both transcription initiation as well as termination sites. The coding region is marked as the larger blue box, which is flanked by the 5' and 3' UTRs indicated by narrower blue boxes. The intronic region is indicated by the thin blue line, with the arrows showing the direction of transcription. Sanger-sequenced PCR products were aligned to the mouse genome GRCm39 (mm39) using minimap2 [1]. Color codes are default for IGV browser [2].

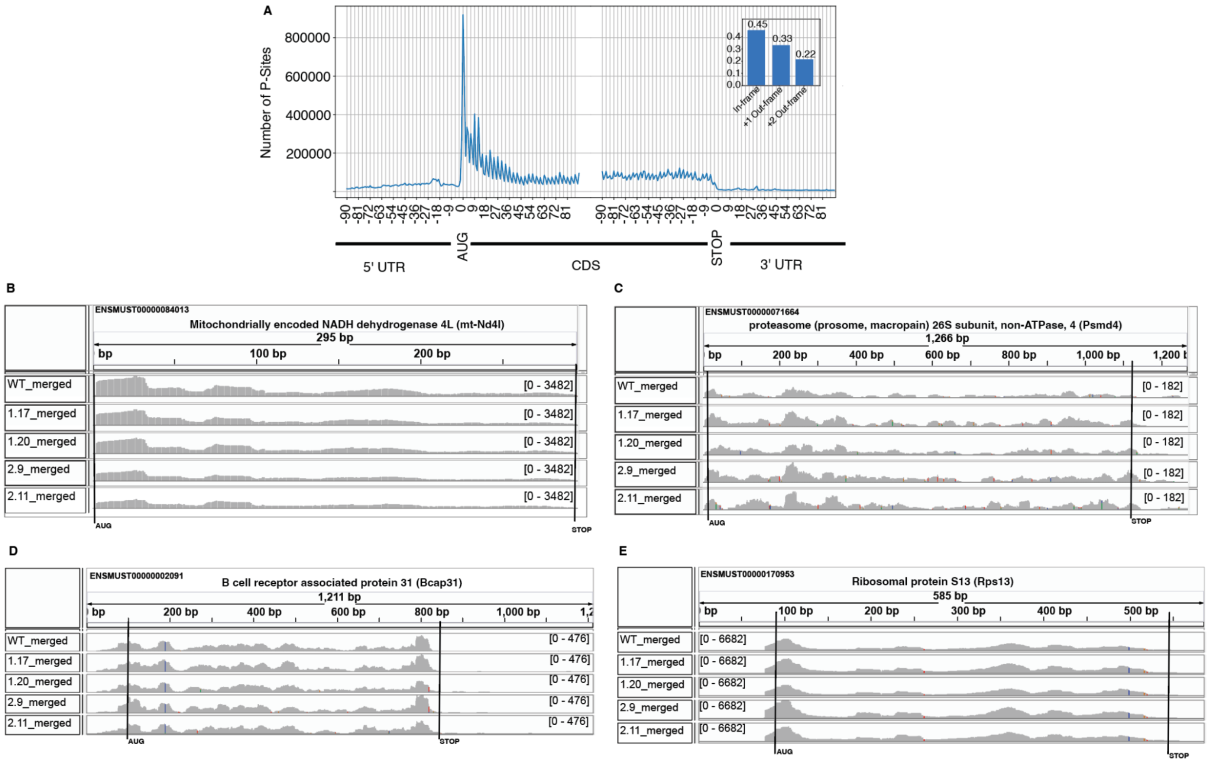


**Figure S3. Quality controls of ribo-seq data from the E14 WT and *Rpl39l* KO clones. A.** Metagene plot showing the frequency distribution of reads mapping to individual positions around translation start (ATG) and stop (STOP) sites. The P sites of decoding ribosomes were inferred from the reads (see Methods) and the numbers of reads supporting each P site were cumulated over all expressed genes. The plot shows the expected depletion of reads upstream of translation start and downstream of translation stop sites. Inset: Proportion of P sites mapping to frame 0, 1, and 2 across all mRNAs. B-E. Integrative genome browser [3] plots showing the coverage of a few transcripts (B - Nd4, C - Psmd4, D - Bacp31, E - Rps13) by ribo-seq data.

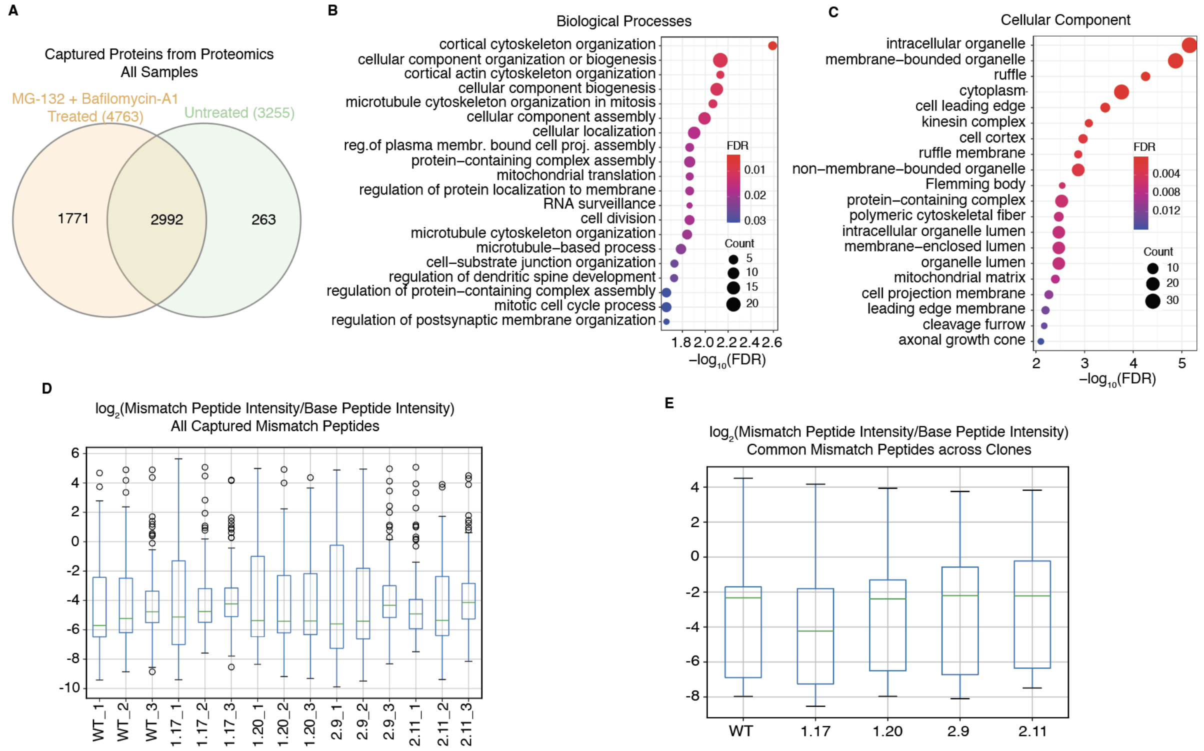


**Figure S4. Analysis of protein stability. A.** Quantification of protein abundance in WT and MG-132+Bafilomycin A-treated E14 cells with global proteomic analysis. **B-C.** Gene Ontology analysis of the proteins whose expression level was significantly downregulated in the *RPL39L* KO clones and restored by the protease inhibitor treatment. Significantly enriched GO terms (x-axis, see also color scale) from the Biological Process (B) and Cellular Component (C) categories are shown. The number of proteins contributing to the enrichment is indicated by the size of the disks. **D.** The mass spectrometry data was used for the analysis of amino acid misincorporation in WT and *Rpl39l* KO E14 cells, with the software developed by Mordret et al.[4]. Base peptides and dependent peptides required for mismatch identification were obtained using MaxQuant computational platform [5] version 2.1.3.0. The analysis was done per clone and per replicate mass spectrometry data set. **E.** Similar to (D), but using only mismatch peptides that were identified in all replicate samples for a given condition.

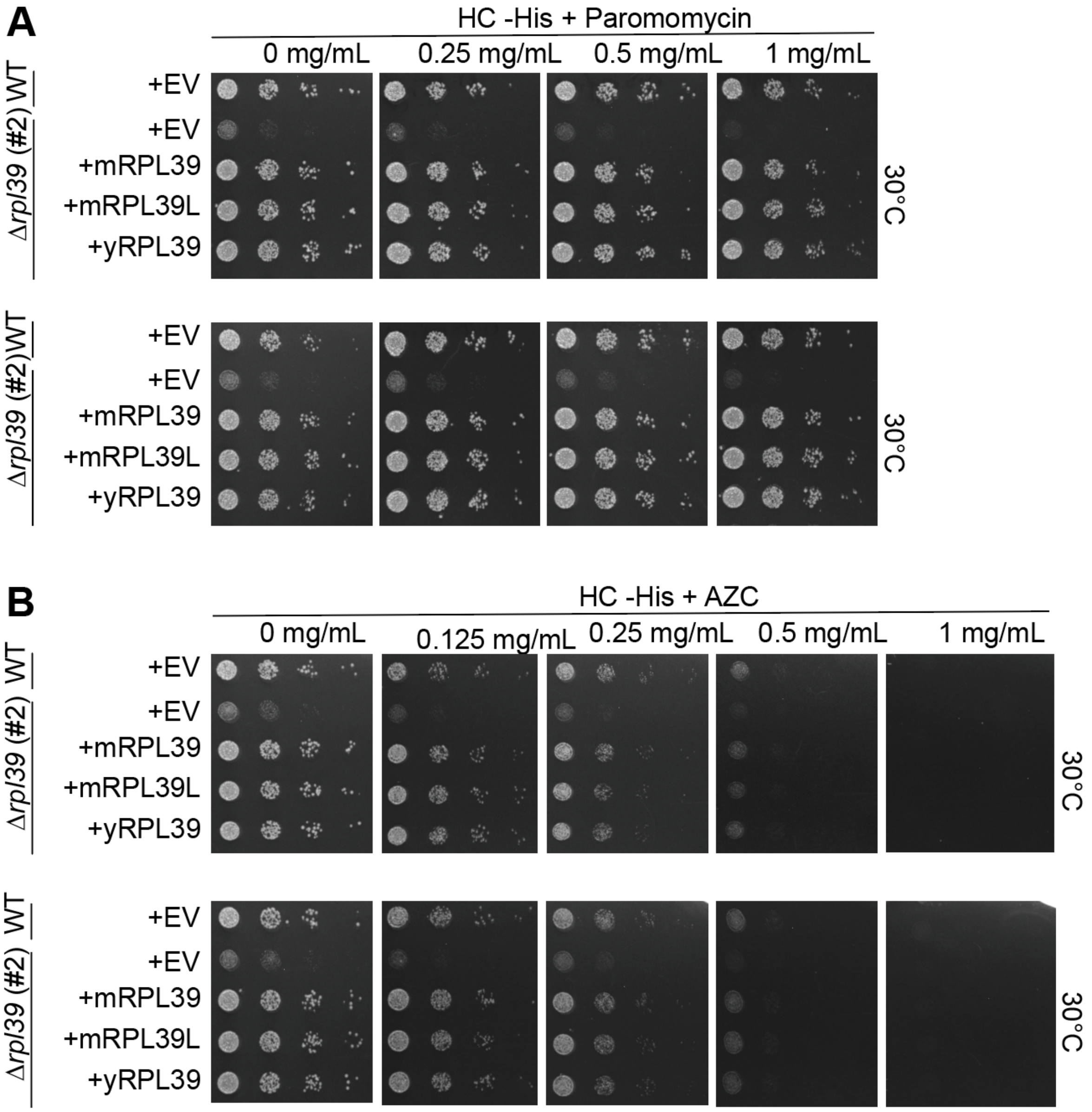


**Figure S5.** ***RPL39*** **KO causes sensitivity to paromomycin and AZC, which is rescued by both mRPL39 and mRPL39L.** The yeast *RPL39* KO lines were transformed with plasmid expressing no protein (empty vector, EV), or the mouse RPL39 (mRPL39) or RPL39L (mRPL39L) of the yeast RPL39 (rescue, yRPL39). EV lines showed strong sensitivity to both paromomycin (A) and AZC (B), and these phenotypes were rescued by the ectopic expression of yRPL39, mRPL39 and mRPL39L.

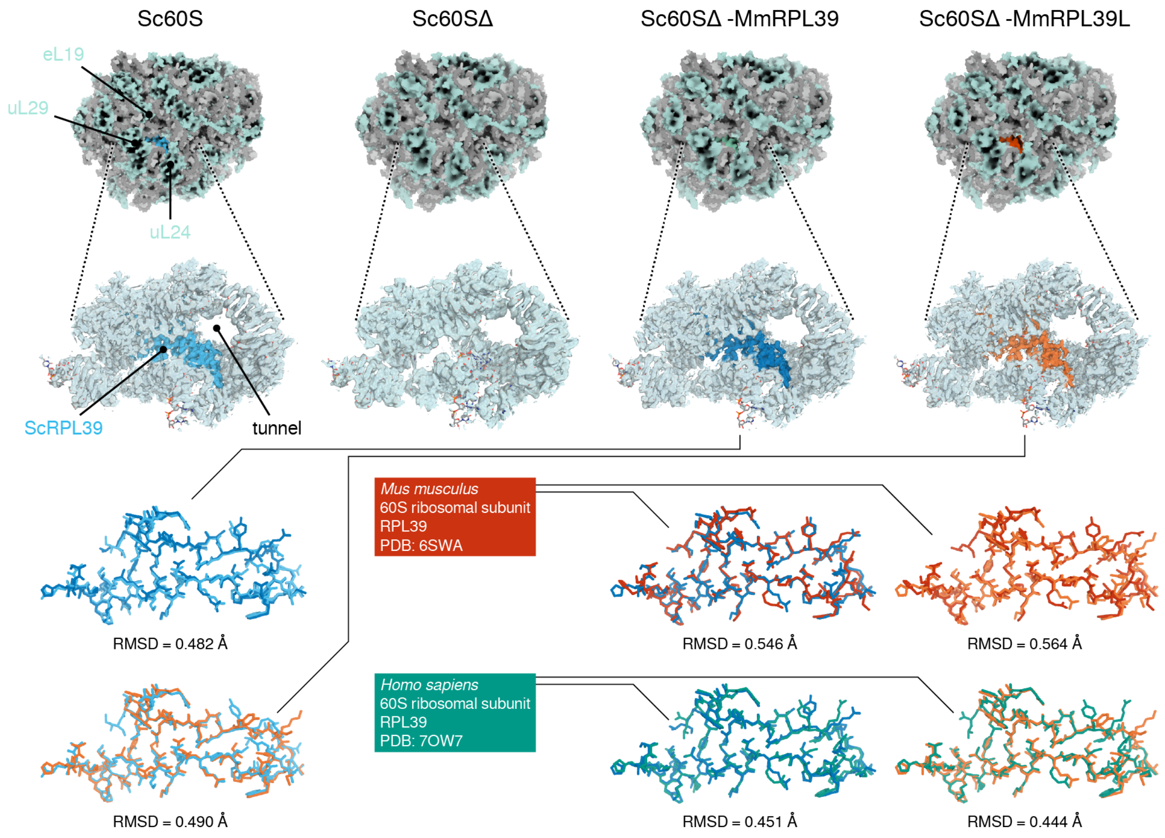


**Figure S6. The structural analysis  of ribosomes containing either mouse RPL39 or RPL39L is possible in heterologous systems.** Ribosomal 60S subunits from RPL39 KO yeast strains with deleted (Sc60SΔ) assemble into intact ribosomal complexes that can be studied by single-particle cryo-EM (top panel, refined atomic model shown in surface representation; rRNA is shown in gray, proteins are shown in pale blue). The position where Rpl39 binds in the WT (shown in cyan) is empty and structural rearrangement in the immediate vicinity is negligible when compared to yeast WT (experimental cryo-EM density shown as pale blue surface, overlaid over refined atomic structure in stick representation). Mouse RPL39 (shown in dark blue) and RPL39L (shown in orange) integrate into ribosomal subunits of the yeast deletion strain, and take the place of the deleted yeast ribosomal protein RPL39. The structures of RPL39 and RPL39Lin the heterologous structures are highly similar to the structures of RPL39 in WT yeast ribosomes and RPL39 in human and mouse ribosomes.

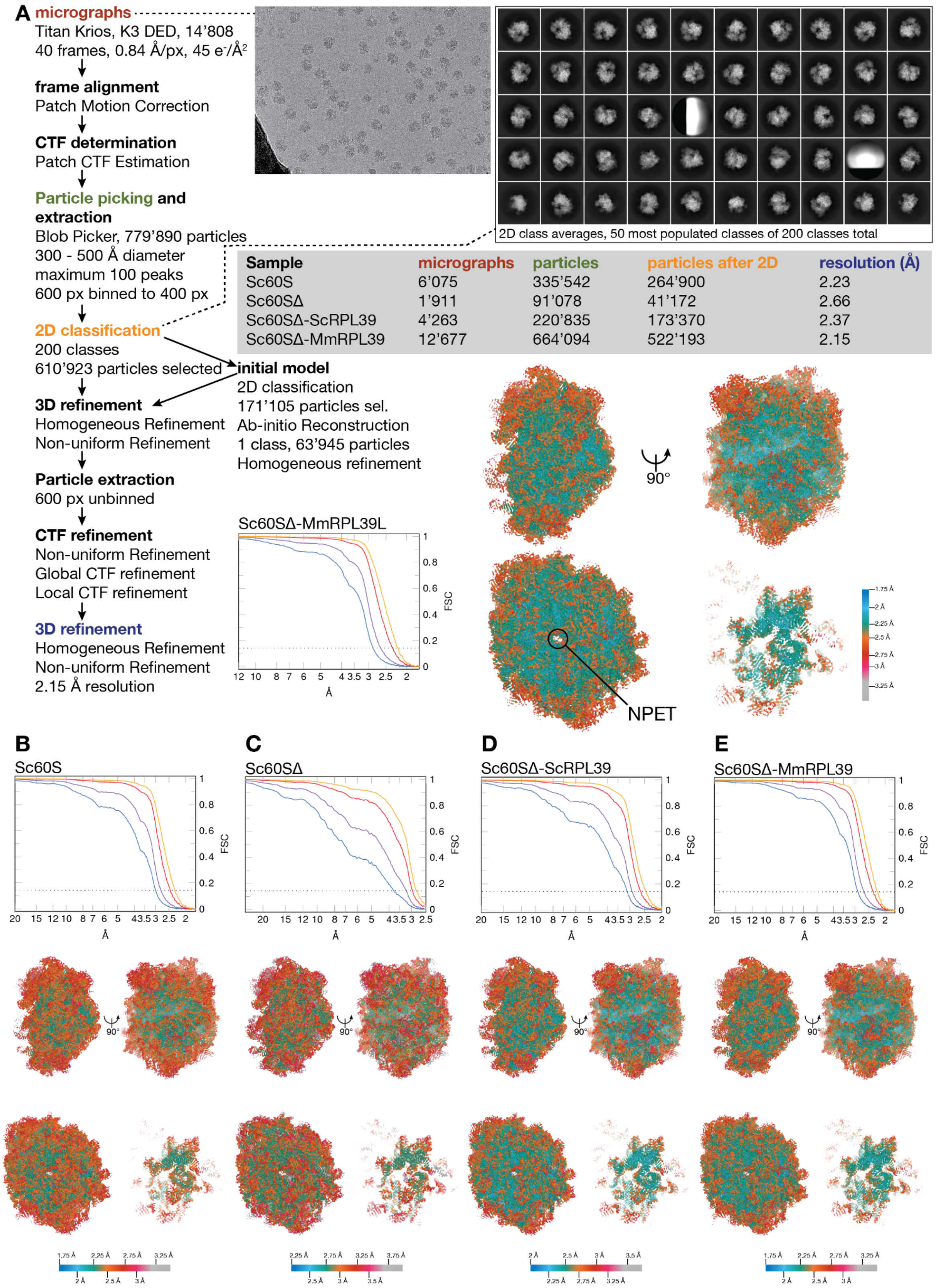


| Sample | micrographs | particles | particles after 2D | resolution (Å) |
|---|---|---|---|---|
| Sc60S | 6'075 | 335'542 | 264'900 | 2.23 |
| Sc60SΔ | 1'911 | 91'078 | 41'172 | 2.66 |
| Sc60SΔ-ScRPL39 | 4'263 | 220'835 | 173'370 | 2.37 |
| Sc60SΔ-MmRPL39 | 12'677 | 664'094 | 522'193 | 2.15 |

**Figure S7. Structure determination by single-particle cryo-EM. A.** Single-particle cryo-EM image processing scheme, Fourier-shell correlation curve, and resolution map of Sc60SΔ-MmRPL39L. Micrograph/particle/particle number after 2D classification, and resolution for the other maps are found in table S5. **B-E.** Fourier-shell correlation curves (calculated with tight/loose/spherical mask in yellow/red/purple, and without masking in blue) and resolution maps of Sc60S (B), Rpl39Δ (C), Rpl39Δ-ScRpl39 (D), and Rpl39Δ-MmRpl39

(E). Resolution maps shown are side view (top left), back view (top right), NPET view (bottom left) and a 25 Å slab around RPL39 residue R28 in the NPET orientation (bottom right).

**Supplementary Tables**

**Table S1. sgRNA sequences used to knock out RPL39L in E14 mESC.**

| **Name** | **Sequence (5'-3')** |
|---|---|
| Rpl39l_ds_1_f | ACACTCAGCTGTCCAGACAG |
| Rpl39l_us_1_f | ACAAAACACAGGGAGTAGGG |
| Rpl39l_ds_2_f | AGTTACAGCATAAGTACGTG |
| Rpl39l_us_2_f | AAACTTGTCCTATGCCATGT |

**Table S2. Antibodies used, concentrations and suppliers.**

| **Antibody** | **Concentration** | **Supplier** |
|---|---|---|
| Eif2alpha Rabbit mAb | 1:1000 | Cell Signaling technology 99722S |
| P-eif2alpha (119A11) Rabbit mAb | 1:1000 | Cell Signaling technology 3597S |
| Perk Rabbit mAb | 1:1000 | Cell Signaling technology C33E10 |
| P-perk Rabbit mAb | 1:1000 | Cell Signaling technology 3197 |
| O-GlcNAc Rabbit mAb | 1:1000 | Thermo Fischer Scientific (MA1-072) |
| Nanog | 1:1000 | Cell Signalling technology 8822 |
| Sox2 | 1:1000 | Cell Signalling Technology 23064 |
| Oct4 Rb pAb | 1:2000 | Abcam Ab18976 Lot: GR3243479-1 |
| Gata4 | 1:250 | Santa Cruz SC-25310 |
| Histone H3 | 1:1000 | Cell signaling 4499S |
| Gapdh | 1:1000 | Santa Cruz SC-32233 |
| Pelo | 1:1000 | Proteintech 10582-AP |
| Vars1 | 1:1000 | Biorbyt ORB373199 |
| Nup93 | 1:200 | SantaCruz sc-374399 |
| Rpn2 | 1:1000 | Proteintech 10576-AP |
| Bcap31 | 1:2000 | Novus Biologicals NBP1-89357 |

**Table S3. gBlock gene fragments used to clone RPL39 mouse and yeast variants.**

| **Name** | **gBlock Sequence** |
|---|---|
| **Mouse RPL39** | 5'- ATG TCT TCT CAC AAG ACT TTC CGA ATC AAG CGA TTC CTG GCC AAG AAA CAA AAG CAA AAT CGC CCT ATT CCT CAG TGG ATC CGG ATG AAA ACT GGT AAC AAA ATC AGG TAC AAC TCT AAG AGA AGA CAC TGG AGG AGA ACG AAG CTG GGT CTG TAA -3' |
| **Mouse RPL39L** | 5'- ATG GCT TCT CAC AAG ACC TTC AGG ATC AAG CGA TTC CTG GCC AAG AAA CAA AAG CAA AAT CGT CCC ATT CCA CAA TGG ATT CAG ATG AAA ACT GGC AAT AAA ATC ATG TAC AAC TCC AAG CGG AGA CAT TGG AGA CGA ACC AAA TTG GGT CTA TAA -3' |
| **Yeast RPL39** | 5'- ATG GCT GCT CAA AAG TCT TTC AGA ATC AAG CAA AAA ATG GCT AAG GCT AAG AAG CAA AAC AGA CCA TTG CCA CAA TGG ATC AGA TTG AGA ACC AAC AAC ACT ATC CGT TAC AAC GCT AAG AGA AGA AAC TGG AGA AGA ACC AAG ATG AAC ATC TAA -3' |
| **Yeast RPL39L** | 5'- ATG GCT GCT CAA AAG TCT TTC AGA ATC AAG CAA AAA ATG GCT AAG GCT AAG AAG CAA AAC AGA CCA TTG CCA CAA TGG ATC CAA TTG AGA ACC AAC AAC ACT ATC CGT TAC AAT GCT AAG AGA AGA AAC TGG AGA AGA ACC AAG ATG AAC ATT TAA -3' |

**Table S4. Primers used to clone Rpl39 variants in yeast.**

| **Name** | **Sequence (5'-3')** | **TM (°C)** |
|---|---|---|
| Fwd_Rpl39_Mouse | GATCTAGAACTAGTGATGTCTTCTCACAAGACTTTCCG | **56** |
| Rev_Rpl39_Mouse | ATGACTCGAGGTCGATTACAGACCCAGCTTCGTTC | **55** |
| Fwd_Rpl39L_Mouse | GATCTAGAACTAGTGATGGCTTCTCACAAGACCTTC | **56** |
| Rev_Rpl39L_Mouse | ATGACTCGAGGTCGATTATAGACCCAATTTGGTTCGTCTC | **56** |
| Fwd_Rpl39_Yeast | GATCTAGAACTAGTGATGGCTGCTCAAAAGTCTTTC | **55** |
| Rev_Rpl39_Yeast | ATGACTCGAGGTCGATTAGATGTTCATCTTGGTTCTTCTCC | **56** |
| Fwd_Rpl39L_Yeast | GATCTAGAACTAGTGATGGCTGCTCAAAAGTCTTTC | **55** |
| Rev_Rpl39L_Yeast | ATGACTCGAGGTCGATTAAATGTTCATCTTGGTTCTTCTCC | **55** |
| Fwd_RPL39_HygroR | ctgaaaattcgaaaaagacaagcaaataaacacagatagatcaacCAGCTGAAGCTTCGTACGC | **57** |
| Rev_RPL39_HygroR | ggaagacaaatgacaaaaagtttgaagcataaatatgttcttcgcGCATAGGCCACTAGTGGATCTG | **57** |
| Fwd_RPL39_Check | ggaagatctggtcatcttggatatg | **56** |
| pAG25R | gagccgtaatttttgcttcg | **54** |

**Table S5: Cryo-EM data collection, processing, and refinement statistics.**

| | **Sc60S** | **Sc60SΔ** | **Sc60SΔ-ScRPL39** | **Sc60SΔ-MmRPL39** | **Sc60SΔ-MmRPL39L** |
|---|---|---|---|---|---|
| **Data collection/processing** | | | | | |
| Magnification | 105’000 | 105’000 | 105’000 | 105’000 | 105’000 |
| Voltage (kV) | 300 | 300 | 300 | 300 | 300 |
| Electron exposure (e–/Å$^2$) | 45 | 45 | 45 | 45 | 45 |
| Defocus range (μm) | -0.5 to -2.5 | -0.5 to -2.5 | -0.5 to -2.5 | -0.5 to -2.5 | -0.5 to -2.5 |
| Pixel size (Å) | 0.84 | 0.84 | 0.84 | 0.84 | 0.84 |
| Symmetry imposed | C1 | C1 | C1 | C1 | C1 |
| Initial particle images (no.) | 335’542 | 91’078 | 220’835 | 664’094 | 779’890 |
| Final particle images (no.) | 264’900 | 41’172 | 173’370 | 522’193 | 610’923 |
| Map resolution (Å) | 2.24 | 2.71 | 2.55 | 2.15 | 2.15 |
| FSC threshold | 0.143 | 0.143 | 0.143 | 0.143 | 0.143 |
| Map resolution range (Å) | 1.854-35.748 | 2.822-46.726 | 1.833-39.904 | 1.854-35.794 | 1.854-35.64 |
| **Refinement** | | | | | |
| Initial model used (PDB code) | 7TOO | 7TOO | 7TOO | 7TOO | 7TOO |
| Model resolution (Å) | 2.4 | 2.9 | 2.6 | 2.4 | 2.3 |
| FSC threshold | 0.5 | 0.5 | 0.5 | 0.5 | 0.5 |
| Map sharpening *B* factor (Å$^2$) | 46.8 | 41.1 | 49.6 | 48.7 | 51.4 |
| Model composition | | | | | |
| Non-hydrogen atoms | 123172 | 124562 | 124998 | 125005 | 125032 |
| Protein residues | 6137 | 6087 | 6137 | 6137 | 6137 |
| Nucleotides | 3388 | 3473 | 3473 | 3473 | 3473 |
| Ligands | Mg: 905 | Mg: 913 | Mg: 913 | Mg: 913 | Mg: 913 |
| | K: 214 | K: 214 | K: 214 | K: 214 | K: 214 |
| | Cl: 70 | Cl: 70 | Cl: 70 | Cl: 70 | Cl: 70 |
| | SPM: 1 | SPM: 1 | SPM: 1 | SPM: 1 | SPM: 1 |

| | | | | | |
|---|---|---|---|---|---|
| | SPD: 1 | SPD: 1 | SPD: 1 | SPD: 1 | SPD: 1 |
| | Zn: 4 | Zn: 4 | Zn: 4 | Zn: 4 | Zn: 4 |
| *B* factors (Å$^2$) | | | | | |
| Protein | 25.53 | 55.46 | 31.63 | 34.36 | 24.63 |
| Nucleotide | 19.48 | 65.14 | 35.65 | 48.83 | 35.52 |
| Ligand | 18.77 | 65.84 | 30.61 | 44.03 | 31.76 |
| R.m.s. deviations | | | | | |
| Bond lengths (Å) | 0.006 | 0.007 | 0.007 | 0.004 | 0.005 |
| Bond angles (°) | 0.729 | 0.690 | 0.749 | 0.664 | 0.697 |
| Validation | | | | | |
| MolProbity score | 1.45 | 1.57 | 1.51 | 1.43 | 1.45 |
| Clashscore | 3.87 | 4.94 | 4.33 | 3.70 | 3.54 |
| Poor rotamers (%) | 0.0 | 0.04 | 0.02 | 0.0 | 0.06 |
| Ramachandran plot | | | | | |
| Favored (%) | 96.02 | 95.52 | 95.70 | 96.02 | 95.59 |
| Allowed (%) | 2.95 | 4.45 | 4.30 | 3.98 | 4.40 |
| Disallowed (%) | 0.03 | 0.03 | 0.0 | 0.0 | 0.02 |
